# The Tensile Expansion Microscopy (TExM) cell stretcher: an iris expansion device integrated with automated real-time autofocus and tracking to super-resolve cells

**DOI:** 10.64898/2026.09.07.749822

**Authors:** Ramita Arampongpun, Tejasvin Shrikanth, Vignesh Venkataramani, Danielle Latham, Marouen Zammali, Vidhi Vakil, Lydia Kisley

**Author notes:** **Correspondence:** Lydia Kisley.

## Abstract

Mechanical stretchers are used to physically expand biological samples for microscopic studies of mechanobiology. Existing stretchers suffer from limited strain capacity and non-quantifiable forces, along with focus drift and feature-tracking limitations during microscopy. We develop an iris-based cell stretcher that applies isotropic equibiaxial force to achieve aerial strain of 1664% that corresponds to 4.2*x* linear expansion for Tensile Expansion Microscopy (TExM), a super-resolution method which increases sample size above the diffraction limit of light. The TExM stretcher uses 3D-printed, printed circuit board (PCB) and cost-effective parts, is portable and automated. We verify the performance of the TExM cell stretcher using image analysis, achieving ∼ 90% mechanical precision, a maximum of four degrees of arm angular deviation, precise speed control down to 0.01 cm/sec, and an average expansion resolution of 3.98 × 10^−3^*x*. Integrated strain gauge sensors confirm equal application of force by each arm of the stretcher throughout expansion and finite-element simulation and planar-polariscope photoelastic imaging characterize the uniform substrate stress distribution while indicating high stress at the substrate gripping area. An AutoTracking software in communication with the stretcher hardware and microscope enables continuous autofocus and feature tracking during TExM and fiducial markers verify uniform equibiaxial stretch. The stretcher is demonstrated with fixed NIH 3T3 fibroblasts and live HeLa cells, observing ∼ 4*x* cellular expansion of fixed cell size and separation of live cell clusters, highlighting the potential of TExM cell stretcher for biological imaging.

**Table of Contents Text:** A 3D-printed, portable, and automated iris-based cell stretcher system integrated with force sensors is developed for Tensile Expansion Microscopy (TExM). The stretcher precisely expands double network hydrogel up to 4.2x linearly. Performance verifications confirm isotropic equibiaxial stretch, 4 degree arm error, and 90% mechanical precision. Applications to track and expand cellular samples show ∼4x fixed fibroblast expansion and live cluster separation.

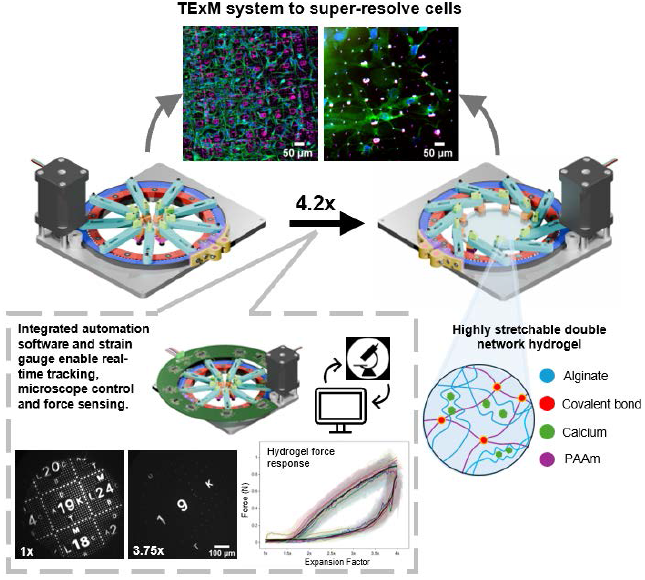

## 1 Introduction

Osmotic Expansion Microscopy (ExM), a super-resolution imaging technique, achieves sub-diffraction s patial resolution on standard microscopes by chemically fixing, hydrogel-conjugating, and enzymatically digesting biological samples to drive isotropic osmotic swelling (4–20 folds, currently up to 1000-fold [1]). This accessible, cost-effective technique is widely used for biological imaging of neural circuits, tumor profiling in microenvironments, and spatial transcriptomics [2]. However, ExM is constrained by a lack of control, reproducibility, and temporal observation. Tissue-variable digestion degrades target fluorophores and causes local structural distortions that require fiducial markers to quantify [3, 4]. Furthermore, ExM requires chemical fixation, preventing live-cell imaging and dynamic temporal studies. Due to the osmotic property, expansion is irreversible and cannot be monitored in real-time under a microscope [5].

Tensile expansion using a mechanical stretcher overcomes the drawbacks of osmotic ExM by physically driving expansion through mechanical force rather than unpredictable fluid uptake. In Tensile Expansion Microscopy (TExM) [5], biological samples are either fixed or adhered to a highly tough and stretchable biocompatible double network hydrogel and subsequently stretched equibiaxially and isotropically in the lateral dimension on an optical microscope [6]. To achieve similar 4x expansion achieved by the initial osmotic ExM studies, aerial strain of more than 1500% is required [5]. The mechanical control of stretching in TExM enables precise, real-time manipulation, dynamic tracking, and reversibility during imaging while avoiding the uncontrolled distortions and restrictive static limitations of osmotic swelling. Therefore, the ability to control the expansion of cells on a deformable material is key to TExM.

Cell stretcher devices have existed for decades to replicate physiological forces (e.g., muscle contraction, digestion, breathing and heartbeat) by deforming the extracellular matrix (ECM), triggering mechanotransduction that regulates cellular functions including proliferation, orientation, reorganization, migration, and apoptosis [7, 8, 9, 10]. The stretcher devices have been developed with balance in strain capacity, dynamic response, biophysical relevance, imaging compatibility, and system complexity [9]. Cell stretchers are categorized into three main configurations based on the direction of forces applied.

Uniaxial stretchers apply a positive strain in the stretching direction and a negative strain in the perpendicular direction [11, 12, 13, 14, 15]. For example, a cam-and-follower mechanism converted rotational motion into 15% linear strain (0-10 Hz) for study of F-actin and focal adhesions, but substrate thinning required extensive manual focusing [12] . Variable-stroke cam-lever-tappet (VSCLT) [13] and stepper motor-driven cell and tissue (CaT) [14] designs achieved higher 20% and 40% strains, respectively, but their excessive weight prevented stage mounting which required a separate ex situ imaging. An Arduino-controlled, 3D-printed microstretcher driven by a bipolar stepper motor and lead screw overcame size, weight, and cost barriers [11, 15], enabling real-time, stage-top imaging, with a future direction stated to integrate a force sensor for concurrent force-deformation tracking.

Biaxial stretchers expand the substrate along two axes and can offer independent directional control with motor actuation or vacuum-driven systems [10]. Pneumatic stretchers such as Flexcell [16] or a dual-channel vacuum device [17] deform elastomeric membranes (silicone, PDMS) around a cylindrical loading post to provide equibiaxial or non-equibiaxial stretch at 15% strain at 1 Hz with <1% error. However, pneumatic latency, strain heterogeneity, focus drift, and substrate viscoelasticity restrict the frequency and strain to <7%. Motor-actuated biaxial stretchers use single-motor indenter rings [18, 19] or dual-motors for independent two-axis control [20] and provide superior strain precision compared to pneumatic devices. However, direct membrane-ring contact introduces friction, resulting in substrate rupture, micro-vibrations, strain heterogeneity, and focus drift. Alternatively, a stage-compatible, software-controlled dual-motor mechanism using a belt-and-pulley system clamped to a free-floating substrate eliminated contact friction achieving strains >200% [20, 21]. However, high stress concentration at the clamps caused the PDMS substrate to rupture prior to reaching maximum stretch.

Isotropic (equibiaxial) stretchers apply symmetric multi-axis forces along six directions, producing lateral expansion coupled with axial compression [22]. Several iris-like stretchers have been developed [23, 24, 25, 26, 27]. The lightweight and microscope-compatible Isostretcher [26] used a rotational swivel mechanism that translated the radial displacement of six hooks to stretch a PDMS membrane, reaching 20% strain within a second with minimal z-drift (∼15 *µ*m). Cells were monitored during low stretch, however, higher strain increased z-drift and risked hook-induced substrate tearing. A similar, yet bulkier system isotropically stretched ∼ 30 mm PDMS substrates up to 20% linear strain via six lip clamps driven by a stepper motor [25]. However, a large shift of focus field (over ∼100 *µ*m) limited cell tracking ability and PDMS autofluorescence interfered with cell stains. A motor-actuated stretcher using eight interdigitating lever arms with mounting pillars screwed to an outer frame allowed translation of rotational displacement to stretch a high extension silicone rubber (HERS) substrate for stem cell culture [24]. The stretcher promised 1000% isotropic surface expansion to study neural tube organoid mechanoregulation on a PEG hydrogel [23], yet, the invasive-designed hooked pillars often cause localized substrate tearing.

The focal shift and resulting out-of-focus fuzzy edges is a major challenge for continuous imaging on higher strain cell stretchers, necessitating custom automated software-based axial autofocus and lateral tracking (AutoTracking). Commercial microscopes use hardware-based autofocusing of the reflection of infrared lasers from static, flat, glass substrates, making it unsuitable for the dynamic experiments with hydrogels for TExM and cell stretching [28, 29]. Passive focusing, an image processing-based autofocus method, relies on gradient-based sharpness evaluation functions (e.g., Tenengrad, Variance, Laplace, Brenner). Slow computational processing, especially in dynamic experiments, have been addressed through combining region-of-interest (ROI) down-sampling, Tenengrad scoring, and Brent search optimization [30]. However, software autofocus alone in the axial direction cannot track cells that drift out of view laterally during high expansion. Comparisons of hardware and software based approaches have been reviewed [31, 32], yet, continuous imaging of cell stretching at the high strains for TExM requires the ability to image the same features over a large field of view and 100’s *µ*m of axial focal change on a soft material.

None of the existing cell stretchers designed for mechanobiology and AutoTracking algorithms offers a combination of high strain (>1500%), compactness and microscope compatibility, low cost, and non-invasive substrate mounting required for TExM. Existing stretcher devices are not feasible to analyze structural dynamics in living cells using high-resolution microscopy during real-time stretching [7, 19]. In addition, quantification of tensile forces have been lacking, as measurements require the integration of a force sensor for real-time quantification during stretch [11]. These ideas prompted us to engineer an advanced and fully equipped stretching device that achieves high strain without tearing the substrate.

Here, we detail our design and verify the performance of an iris-based electromechanical stretcher for tensile expansion microscopy (TExM) (Figure 1). The system delivers uniform equibiaxial stretching at an extreme expansion factor up to 4.2*x* (1664% strain) without the extensible double network hydrogel substrate tearing, superseding the strain limits of existing cell stretchers. The stretcher is engineered with a user-centered design for ease of use, standard microscope compatibility, low manufacturing cost using 3D-printed parts, long operational lifetime, and an efficient sample loading procedure. Our TExM stretcher provides precise control over stretching parameters such as speed and stretching step size. Embedded strain gauges enable real-time continuous strain measurements and we confirm equibiaxial expansion and precision of the stretcher through finite element simulation and experimental analysis. The AutoTracking software integrated with the hardware resolves focus drift and allows tracking of the same cell or two-photon lithography fiducial markers to streamline high-resolution imaging workflows. Finally, we apply the stretcher to expand fixed and live cells demonstrating increased resolution for structural subcellular or single cell analyses, respectively.

**FIGURE 1.**
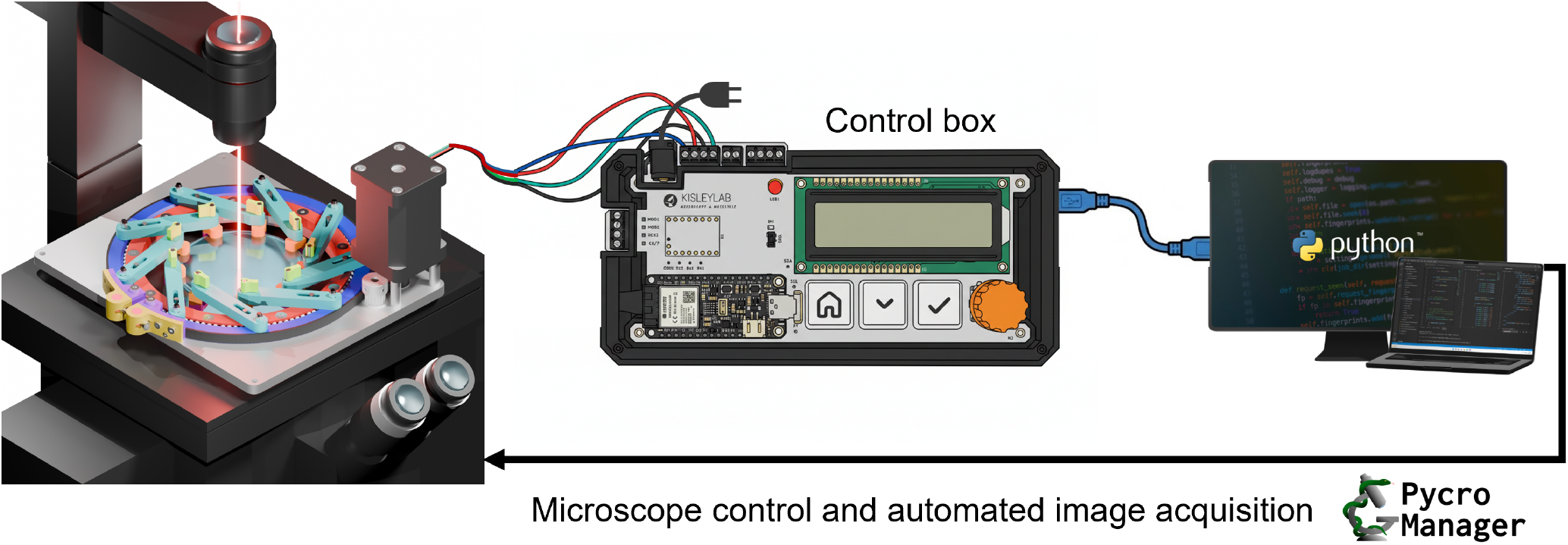
Overview of the TExM stretcher system

## 2 Description of the TExM Stretcher System

### 2.1 Stretcher Device Design, Justification, and Implementation

The cost-effective TExM system is designed to use approachable 3D-printed parts and expand samples based on an iris stretcher design (Figure 2). Figure 2a shows the details consisting of eight main parts: a base plate, rings, nylon beads, belt and pulley system, stepper motor, arm, and gripper. Here, we describe each component and the justification for the design choices made in relation to our application to expand cellular samples for optical microscopy.

**FIGURE 2.**
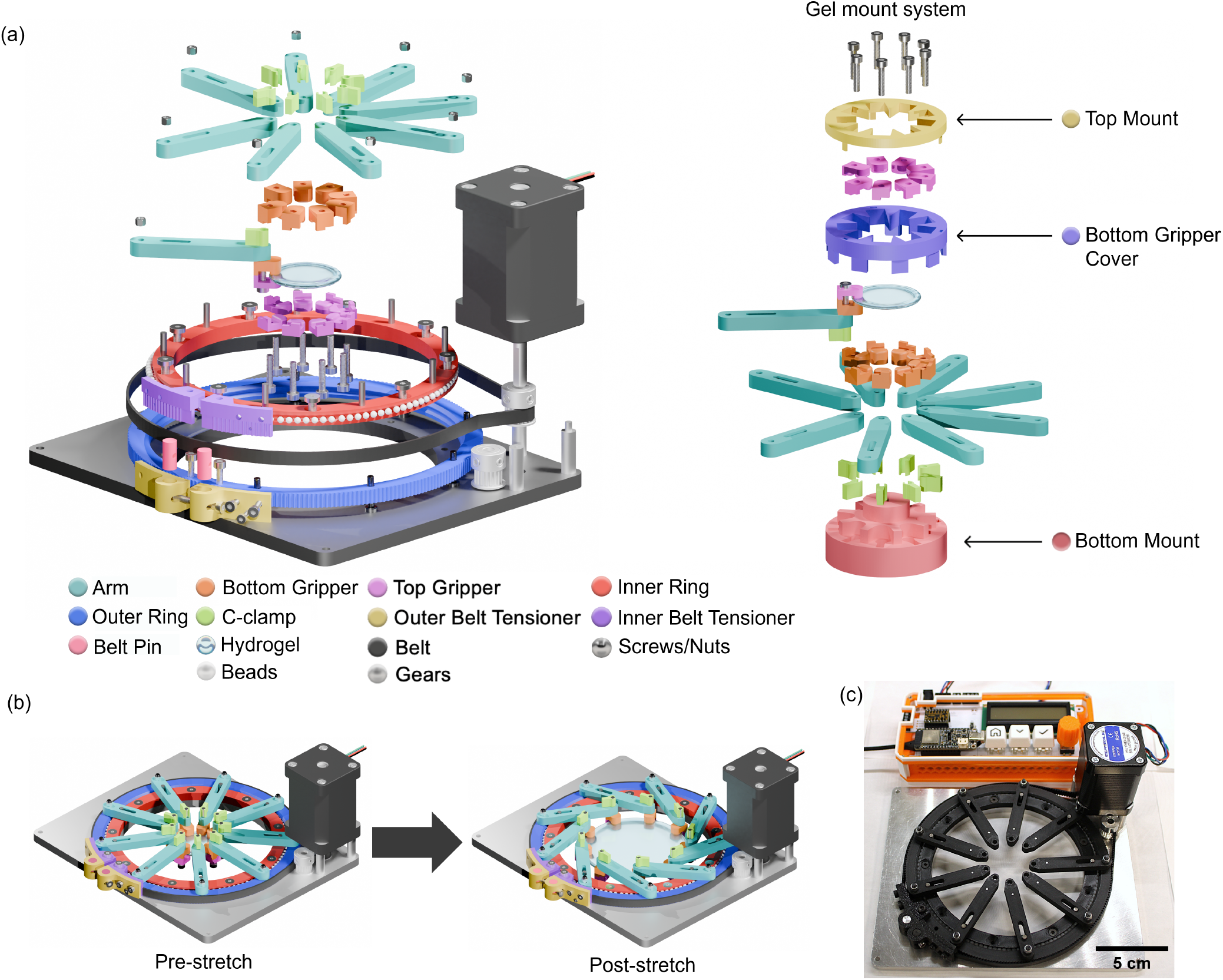
All parts of TExM stretcher system. (a) Exploded view of TExM stretcher device and sample loading mechanism, (b) Demonstration of a hydrogel during pre- and post-stretch on TExM stretcher, and (c) Photo of stretcher prototype with the control box in orange.

Materials were selected for weight, strength, cost, accessibility, and prototyping. The base plate is made of aluminum sheet (89015K324, McMaster Carr) cut using a CNC machine and is designed to fit on a standard microscope stage for imaging purposes. The size allows for the stretcher to be “dropped in” for ready use on an Olympus microscope, but also limits the achievable expansion factor based on the geometry. Aluminum has strong rigidity compared to acrylic, corrosion resistance to any biological buffers, and is light given the 1 kg weight limitation of the microscope translation stage. All other parts are 3D printed using a Bambu Lab H2S printer with polylactic acid (PLA) and acrylonitrile butadiene styrene (ABS) filament materials for fast prototyping, low cost, ease of assembly and replacement, while offering good tensile strength and chemical resistance properties.

The belt and pulley system, nylon beads, rings, and bipolar stepper motor (Nema 17 model 17HS24-2104S, StepperOnline) are used to ensure smooth rotation of the stretcher during expansion. An inner ring is fixed to the base plate while an outer ring rotates freely with the belt. Nylon beads of 3.18 mm in diameter (9613K13, McMaster-Carr) act as ball bearings to reduce friction between the two rings. Nylon is selected for high strength, corrosion and wear resistance, and self-lubrication. The timing belt (3682N1, McMaster-Carr) is wrapped around the stepper motor pulley (3684N12, McMaster-Carr) and the outer ring, transferring tension forces to move the arms. There is one idle pulley for supporting the belt and dissipating forces acting on the motor pulley. Belt tensioners are used to secure and connect both ends of the belt by three screws, two M2 screws fix the belt to its tensioners while the other four M3 screws help support the attachment and reduce movement by sandwiching the belt between the inner and outer tensioners. The multiple screws and tensioners hold the belt in place without slipping or slacking, allowing for tunable tightness when expanding a wide range of substrate stiffnesses (Figure S1 detailed a post-assembly image). Further, the stepper motor is secured to the base plate using three M3 screws on three aluminum bar standoffs (8252T511, McMaster-Carr), which are cut and threaded to match the height of the motor. The motor is controlled by the driver (TMC2209, DigiKey) on the control box.

Multiple arms and rotatable, rough, friction-modified grippers [33] hold the hydrogel in place and dissipate force. Nine arms are attached to the outer ring. The large number of contact points dissipate force around the circumference of the pre-expansion hydrogel; nine, while not ideal for symmetry, was the maximum contact points that could fit in the pre-expansion state (Figure 2b). The end of the arms are in contact with the hydrogel through a c-clamp-based gripper. The grippers consist of a top and bottom portion, attached on either side of the arm via a c-clamp and screw to allow for self-rotation when resistance force of the hydrogel is present during expansion. Alternative approaches using a nut limited rotation, resulting in an inconsistent distribution of torque hydrogels tearing, and anisotropically expansion due to shear, causing obvious distorted images of biological samples. The c-clamp grippers further require high surface friction to hold the hydrogel in place during expansion achieved by bonding P60 grit sandpaper (50-10045, Allied High Tech Products Inc.) with adhesive (75445A44, McMaster-Carr). In the absence of sandpaper, the hydrogels slip out of the grippers, even with mm-height sawtooth texturing. Alternatively, physically screwing through the hydrogel introduces tears and creates starting sites of gel failure. Finally, grippers need as small dimensions as possible (25.67 mm × 7.94 mm) to achieve the maximum expansion factor and imageable surface area while successfully expanding hydrogel without slipping or tearing.

The post-expansion size of hydrogel is related to the gripper area and overall stretcher parts dimensions which reflects the maximum achievable expansion factor (Figure S2). The smaller the pre-expansion hydrogel, the larger expansion factor can be achieved based on the limitation on the base plate size. With the constraints to grip the edge of the hydrogel with the nine arms and the stretcher geometry, the smallest starting hydrogel diameter is 30 mm. This further corresponds approximately to the size of a standard six-well cell culture plate (diameter = 35 mm), where the 35 mm size can be used-as is for lower expansion factors or cut to size. Figure S2 compares between the two designs of the grippers and mounting system affecting the maximum expansion factor.

A 3D printed gel mount system made of PLA plastic assists load the sample onto the stretcher (Figure 2a). The hydrogels are soft, flexible, and contain sensitive biological samples. Lifting and transferring actions in the absence of a support poses high risk for damaging the sample; cells can fall off of the surface and difficult handling by the user can lead to physical removal of cells or non-controlled expansion of the hydrogel, breaking double network bonds. Figure S2 illustrates different versions of gel mount system. Loading the hydrogel sample using the gel mount system starts with placing all c-clamps into their slots. The bottom mount is designed to have slots for the arms to perfectly fit by overlaying the arms on top of the bottom mount. After assembling the bottom mount with the arms, the bottom gripper cover is placed over the arms from the top which serves as holding slots for the bottom grippers and prevents the grippers from moving while loading the samples. Once the bottom gripper cover is in place, the bottom grippers are arranged into the slots. The hydrogel sample is placed into the center and the top mount is placed to guide attachment of the top grippers with M3 screws. The top mount has small legs which fit to the slot on the bottom gripper cover to prevent slipping or misalignment while securing the hydrogel. Once the hydrogel is assembled, c-clamps secure and prevent the hydrogel from falling or lifting up; all parts of the gel mount system can be easily lifted off while leaving the hydrogel clamped to arms of the stretcher.

The stretcher is operated by a user-friendly, compact, and modular control box. A printed circuit board (PCB) designed in KiCad and manufactured by JLCPCB (Figure 1 and 2c) with controller integration is used with the rotary encoder (PEC11R-4220F-S0024, DigiKey), switches (MX2A-E1NA, DigiKey), and liquid crystal display (LCD) (NHD-0216K1Z-FSW-FBW-L, DigiKey). Apart from being light weight and portable, the high modularity of the PCB control box reduces noise and makes troubleshooting easier compared to wired breadboards (Figure S3). Furthermore, it supports future modifications for additional functionality such as wireless control through ESP32 Wi-Fi antenna. Most importantly, the PCB schematic is sharable and easy to manufacture due to less expertise required, thus increasing the reproducibility of the part.

After the hydrogel is gripped and loaded onto the stretcher using the sample loader, the nine arms are moved radially to generate an equibiaxial stretch to the gripped hydrogel (Figure 2b) as a result of belt system movement caused by a stepper motor controlled through a microcontroller (Adafruit ESP32-S3, DigiKey) on the PCB board. Figure 2c shows a photo of the prototyped, constructed device. Additional 3D printed parts are shown in (Figure S4). Earlier prototypes and discussion of the grippers, gel mounting system, and control box are provided in the Supporting information (Figure S1-4). An instructional assembly video and stretcher manual are available at Kisley lab Github .

### 2.2 TExM Stretcher Control

The TExM stretcher can be controlled in two ways: manually on the control box (Figure 2c) and software-controlled through Python for automated tasks (described in section 4). The controllable parameters, such as stretching speed and step, can be adjusted through a custom C++ based code run via Arduino software. The code consists of five main functions: the “SetZero”, which happens once the device is connected to the PC, will acquire the current position of the arms and set it as a zero. Before running the “GoTo” function to move the iris arms to the desired expansion based on the equation 3 in Section 2.3, users can select the desired speed using the ‘Speed’ function which controls the rate of the arms movement in cm/sec. The “GoTo” function can also be used to return the hydrogel to lower expansion. The “Zero” function moves the arms to the initial zero position saved. The “Calibrate” function is beneficial for resetting the zero and max position when the stretcher is being used continuously without disconnecting. It works by automatically expanding the arms in both clockwise and counter-clockwise directions to identify the maximum expansion from both directions for recalculating the zero position.

### 2.3 Derivation of the 4.2*x* Expansion Factor Based on Stretcher Geometry

The expansion factor is derived from a geometric calculation of the angular movement of the TExM stretcher arms relative to stepper motor counts. As the motor moves the belt, the outer ring rotates which turns the arms outward at a defined angle until reaching the maximum travel distance when the arms touch the inner ring. The schematic diagram of the arm’s geometry is shown in Figure 3a.

**FIGURE 3.**
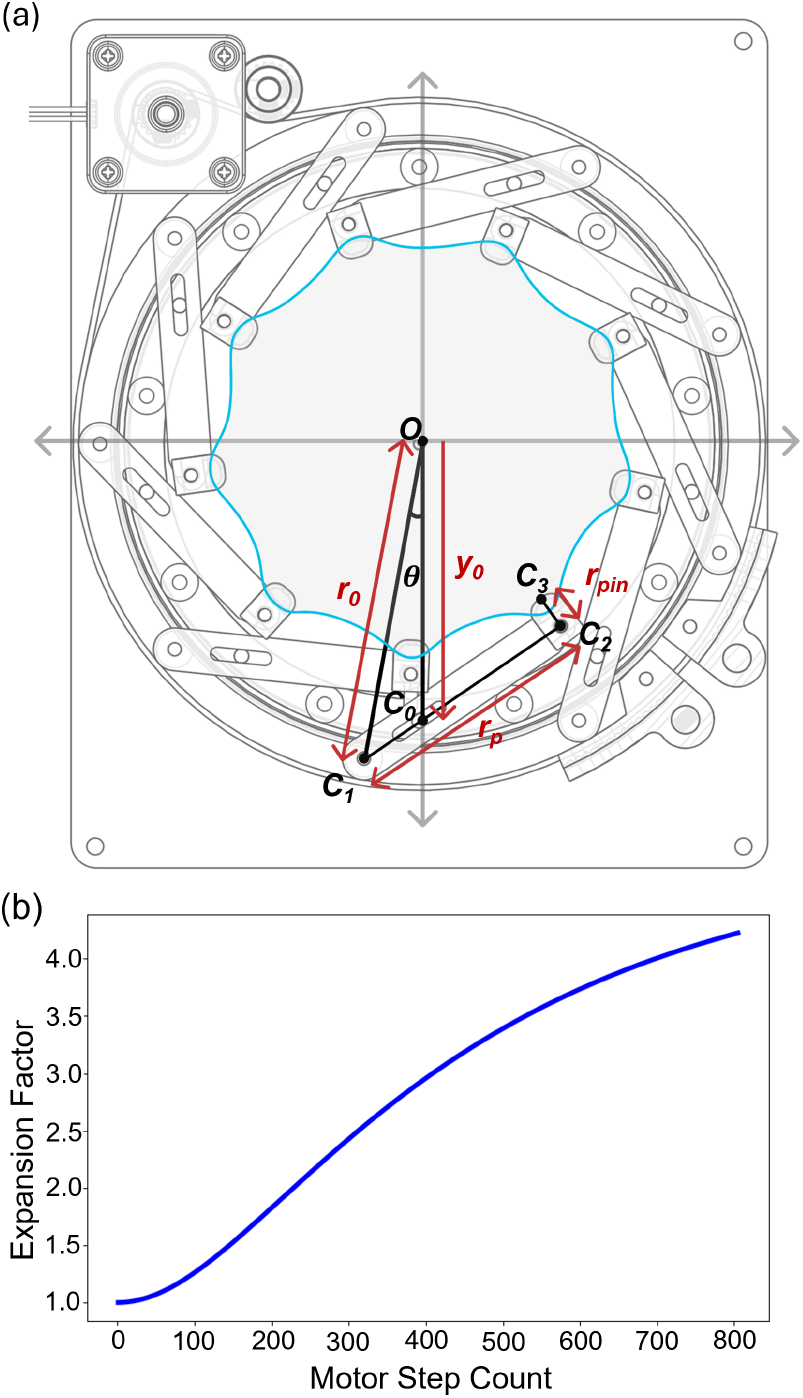
TExM stretcher mechanism kinematics and components. (a) Schematic of the expansion of the hydrogel with geometric variables defined for one TExM arm at its maximum expansion. (b) Expansion factor and motor step count derived from Equation 3

The device’s gear ratio (g) dictates the proportional angular displacement of the outer ring relative to the stepper motor input which can be derived as g = *r*_*rin*g_ /*r*_g*ear*_ where *r*_g*ear*_ is the motor pulley radius and *r*_*rin*g_ is the outer ring radius. The relationship between the stepper motor angle and the angle of the outer ring is *θ* = *θ*_*motor*_ /g where *θ* is the outer ring angle (degrees), *θ*_*motor*_ is the stepper motor angle (degrees), and g is the gear ratio. Microstepping enables smooth, high-resolution movement of the stretcher arms. Equation 1 defines the motor pulley angle, where *N*_*steps*_ = 200 (full steps per motor revolution), *N*_*micro*_ = 200 (maximum microstepping factor of the motor), and *S* is the number of microsteps moved.

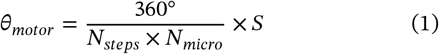

The relationship between the angle of the outer ring and the expansion factor can be obtained by determining constants, variables, and points on the stretcher (Figure 3a). The constant parameters include radius from the center *O* to the outer ring *c*_1_ which connects to the end of arm (*r*_*o*_), length of the arm (*r*_*p*_), distance from the fixed pin on the inner ring at point to the center (*E*_*o*_ = *r*_*o*_ − *r*_*p*_), and the distance between the tip of the arm to the center (*θ*). Variables include the angular position of the outer ring (*θ*) and the angular position of the stepper motor (*θ*_*motor*_). Other points include the opposing end of the arm which connect to the grippers (*c*_2_) and the tip of the gripper (*c*_3_). Assuming the points have the following coordinates: *c*_0_ = (0, *y*_0_), *c*_1_ = (*x*_1_, *y*_1_), *c*_2_ = (*x*_2_, *y*_2_), and *c*_3_ = (*x*_3_, *y*_3_) then, *x*_1_ = *r*_0_*cos*(*θ*) and *y*_1_ = *r*_0_*sin*(*θ*). Using the unit circle equation stating *x*_1_^2^ + *y*_1_^2^ = *r*_0_^2^ and the distance between the points *c*_1_ and *c*_2_ is a constant equal to the length of the arm (*r*_*p*_), we get *r*_*p*_ ^2^ = (*x*_1_ − *x*_2_)^2^ + (*y*_1_ − *y*_2_)^2^. To obtain *x*_2_ and *y*_2_ in terms of *θ*, we know that *c*_2_ is on a straight line connecting *c*_1_ and *c*_0_, thus 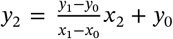 and *x*_2_ can be derived from quadratic equation as:

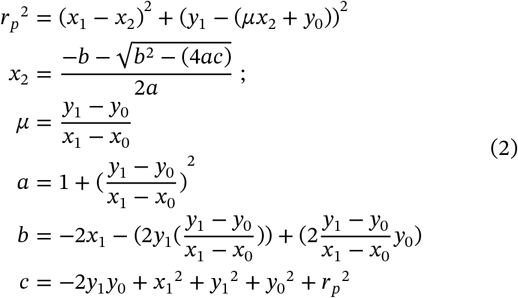

Finally, the expansion factor can be calculated by the distance from the point to the center divided by the pre-stretch state of the hydrogel (relaxed state) in Equation 3. Note that the gripper is pointed towards the center due to the tension of the hydrogel.

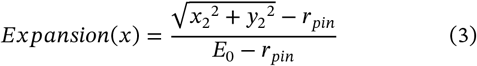

Using the constant dimensions of g = 12.7, *r*_0_ = 7.25 cm, *r*_*p*_ = 5.32 cm, *y*_0_ = -6.19 cm, and *r*_*p*_ = 0.766 cm of the TExM stretcher design results in the relationship between expansion factor and stepper motor count in Figure 3b.

The stretcher geometry yields maximum linear expansion of 4.2*x* with speed ranges from 0.01 to 5 cm/sec. At the minimum actuation speed of 0.01 cm/second, complete expansion requires approximately 33 seconds. Rotational-to-linear motor motion produces a sigmoidal-like displacement profile (Figure 3b). There are three distinct expansion zones: initial zone (∼1-150 steps) where expansion starts off slowly, linear region (∼151-500 steps), and saturation zone (∼501-800 steps) where rate of expansion slows down as it approaches maximum limit. Due to the non-linear nature, average expansion factor resolutions per step of each zone are 3.43 × 10^−3^*x*, 5.27 × 10^−3^*x*, and 2.76 × 10^−3^*x*, respectively. Overall, the stretcher demonstrated high positional fidelity, maintaining a mean mechanical resolution of 3.98 × 10^−3^ expansion factor (*x*) per step.

## 3 Performance Verification of the TExM Stretcher

### 3.1 Engineering Performance and Angular Tolerance

The TExM stretcher arm tolerances quantify the performance and precision when stretching a sample. We compare the engineering performance on two states of the stretcher: a control state without any tensile force acting on the arms and a standard sample operating state which includes a hydrogel (SI section 2 described hydrogel preparation method) mounted on the arms using the grippers. Quantification was done through image analysis using circular markers on the TExM stretcher’s arms. In the control state, the void screw holes in the absence of the grippers at the end of the arms were used (Figure 4a, inset, yellow circle), as the grippers would freely rotate without the tension of the hydrogel. For the standard operating state circular markers made of white tape cut using a hole puncher, were placed at the end of the grippers (Figure 4b, inset, yellow circle). The voids or white markers provide high contrast with the black arms for image binarization. Despite the marker size and location differences, the angular measurements of control and sample states are consistent and do not affect the analysis. Stepwise expansion videos were recorded using a Canon digital camera (EOS Rebel T7 with EF-S 18-55 mm f3.5-5.6) placed above the TExM stretcher using a tripod and obtained a top-down view of the expansion at 30 fps. A LED white light illumination pad (Huion L4S light pad 1100 Lux, Amazon) was placed behind the stretcher to eliminate any shadows. Images were pre-processed using ImageJ software by applying edge finding and binarization via auto thresholding then manually inspected to remove artifacts from lighting from the binarized image.

**FIGURE 4.**
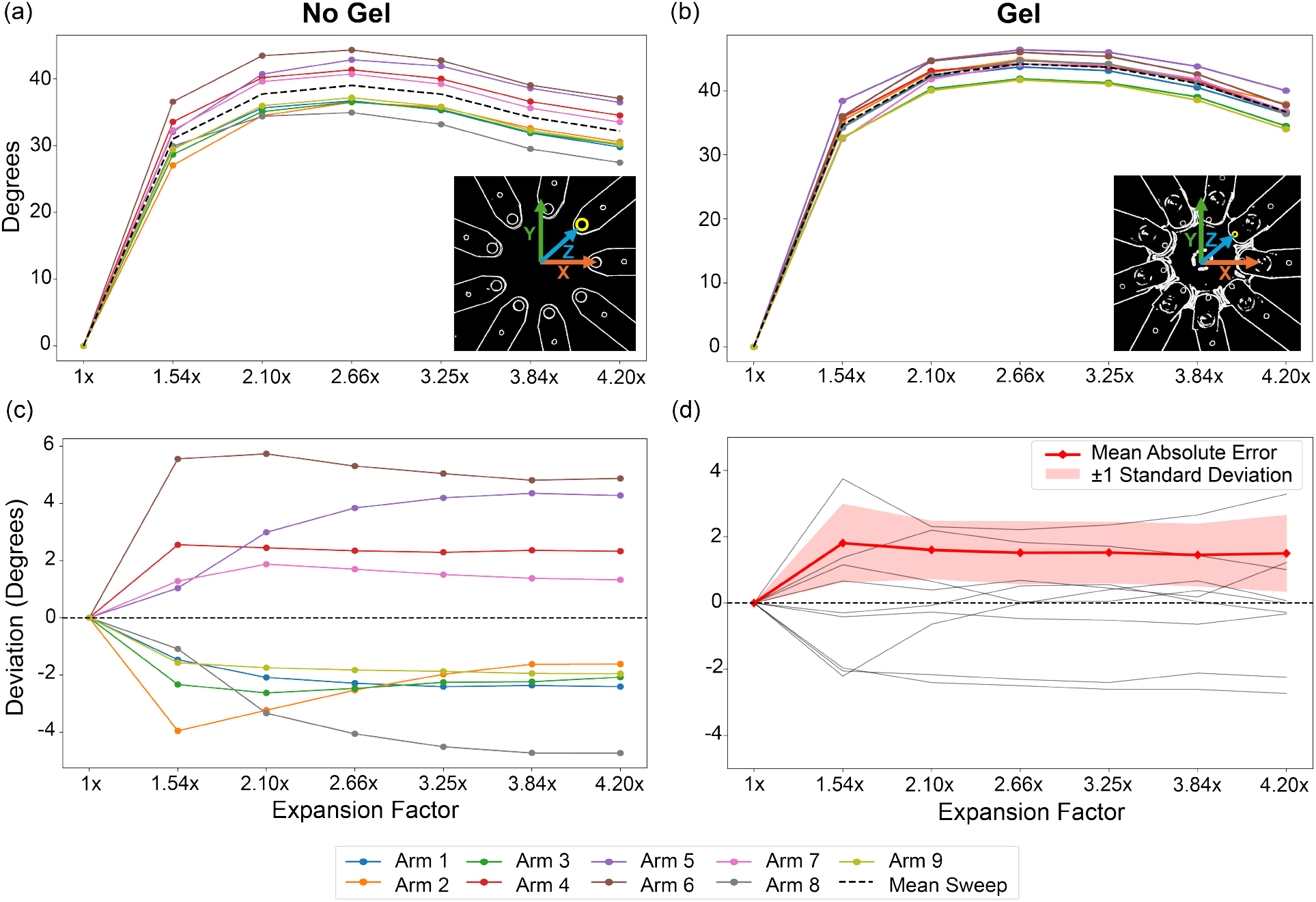
Quantitative analysis on TExM stretcher’s angular tolerance of nine arms individually on control state (a, c) where there is no hydrogel mounted and standard sample state (b, d) with presence of hydrogel. One control and three samples (standard state) were analyzed. The inset images show binarized images used for analysis. (a, b) illustrates angular displacement in degrees of each arm in relation to expansion factor while (c, d) shows the deviation error of each arm from the mean at specific expansion factor. The deviation error plot in (d) is generated from three hydrogel samples, the faded line is deviation error from each arm while the red line represents absolute mean error at each expansion factor and its standard error from the three samples.

Binarized images were analyzed using Python and OpenCV libraries [34] to determine the angular deviation between arms on the stretcher. The circular markers in post-processed images were identified based on size exclusion (area) and circularity. The center point of the stretcher (*x*_*c*_, *y*_*c*_) was quantified relative to all nine markers:

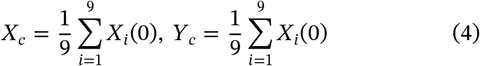

Directly identifying the center point from the markers reduces variability from human error produced during arms assembly and inter-sample differences of the initial direction of grippers when the hydrogel was mounted. After obtaining the center point from the pre-stretch image (1*x* expansion), the marker identification process was repeated for all other images at each controlled stepwise expansion. According to Equation 3, changes in angular movement are related to the change in expansion factor. Thus, each arm’s angular movement (*θ*_*i*_) can be computed as:

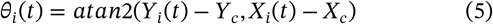

The angular displacement of individual arms at each expansion can be determined by subtracting the angular movement (*θ*_*i*_) at the current expansion from the arm’s angle at initial position (prestretch).

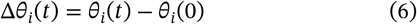

Deviation error (*δθ*_*i*_ (*t*)) can then be calculated by subtracting the actual angular displacement from the mean displacement of all nine arms, represented as:

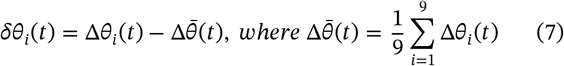

Mechanical precision (*MP*) is a measurement of how the stretcher performs if assuming an ideal operation without any arm’s deviation means the precision is at 100%. After obtaining maximum value from the deviation error, the TExM mechanical precision can be computed as:

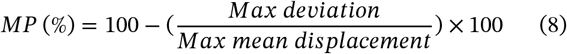

The TExM stretcher successfully demonstrates the high engineering performance in arms tolerance for consistency in tensile force application to expand the substrate. Figure 4 shows the quantitative analysis of TExM stretcher arms tolerance where control and sample deviation error are compared at different expansion factors. The TExM stretcher operates with mechanical precision of ∼ 90% and has a maximum of 4° arms deviation error (absolute average ∼ 2°) when stretching hydrogels.

The TExM stretcher has non-linear changes in angle during expansion with a similar trend of the stretcher arms deviation error between control and sample. Based on the stretcher design (Section 2.1), there is a large increase in arm’s angular displacement at 1.54*x* then gradual increase to reach a peak displacement at 2.66*x* before a slight decrease when it reaches a full expansion (Figure 4a, b). The control has the maximum mean angular displacement of 39°, the maximum deviation error of 6°, and mechanical precision at 85.3%. The sample has the maximum mean angular displacement of 44°, the maximum deviation error of 4°, and mechanical precision at 90.2%. Due to hydrogel resistance force acting on the gripper during expansion, the sample has slightly higher arm angular displacement and lower in deviation error compared to the control due to more stability introduced to the arms. A Welch *t*-test (unequal variance) revealed no statistically significant difference in arm deviation error between control and sample conditions (*p* = 1.00). Thus, individual arm angular displacement remained consistent across both test conditions regardless of hydrogel presence, demonstrating reliable operation of the stretcher unaffected by assembly tolerances in the 3D-printed components, base plate, or rings.

### 3.2 Tensile Force Characterization Using Strain Gauges

Force sensors using strain gauges are integrated for real-time measurement of tensile force exerted on the hydrogel on the TExM stretcher. Stress-strain profiles of hydrogels are well understood under uniaxial loading, translating to two-dimensional equibiaxial conditions is less established due to mechanical tolerances and macroscale deviation from purely isotropic expansion in which the strain gauge can characterize [35, 36]. Linear strain gauges have been used as force sensors in biomedical applications [37]. The strain gauge is a thin, patterned conductor (usually made up of foil or sputtered metal film) bonded to an insulating backing. When a substrate attached to it stretches or compresses, the gauge length and cross-section change, altering its electrical resistance. By analyzing how the hydrogel responds to uniform in-plane stretching, insights into rheological properties and mechanical behavior, such as elasticity, nonlinearity, and time-dependent deformation can be determined, enabling more accurate modeling of hydrogel behavior in the context of TExM.

#### 3.2.1 Strain Gauge Assembly & Procedure

Since the relationship between the applied force on the hydrogel and resulting expansion is not directly measurable, we treat the stretcher as a load cell, mounting strain gauges at various points on the stretcher arms to measure deformation during hydrogel expansion (loading) and contraction (unloading). A total of eighteen strain gauges were used, two per arm across the nine arms, wiring both compression and tension together in a half-bridge arrangement to resolve twice the force sensitivity for a given arm (Figure 5a). The two gauges on each single arm were wired in series on a custom PCB across a Wheatstone bridge in the half-bridge configuration (SI section 3.1). This doubles the bridge output through the opposing resistance changes and cancels common-mode temperature drift. Data collection was driven by Arduino code interfacing the iris stretcher with the PCB, which streamed the step count, ADC values, and timestamp over the serial bus to be recorded on an ESP32 microchip; a Python serial reader using [38] logged this stream and exported it to CSV files for analysis.

**FIGURE 5.**
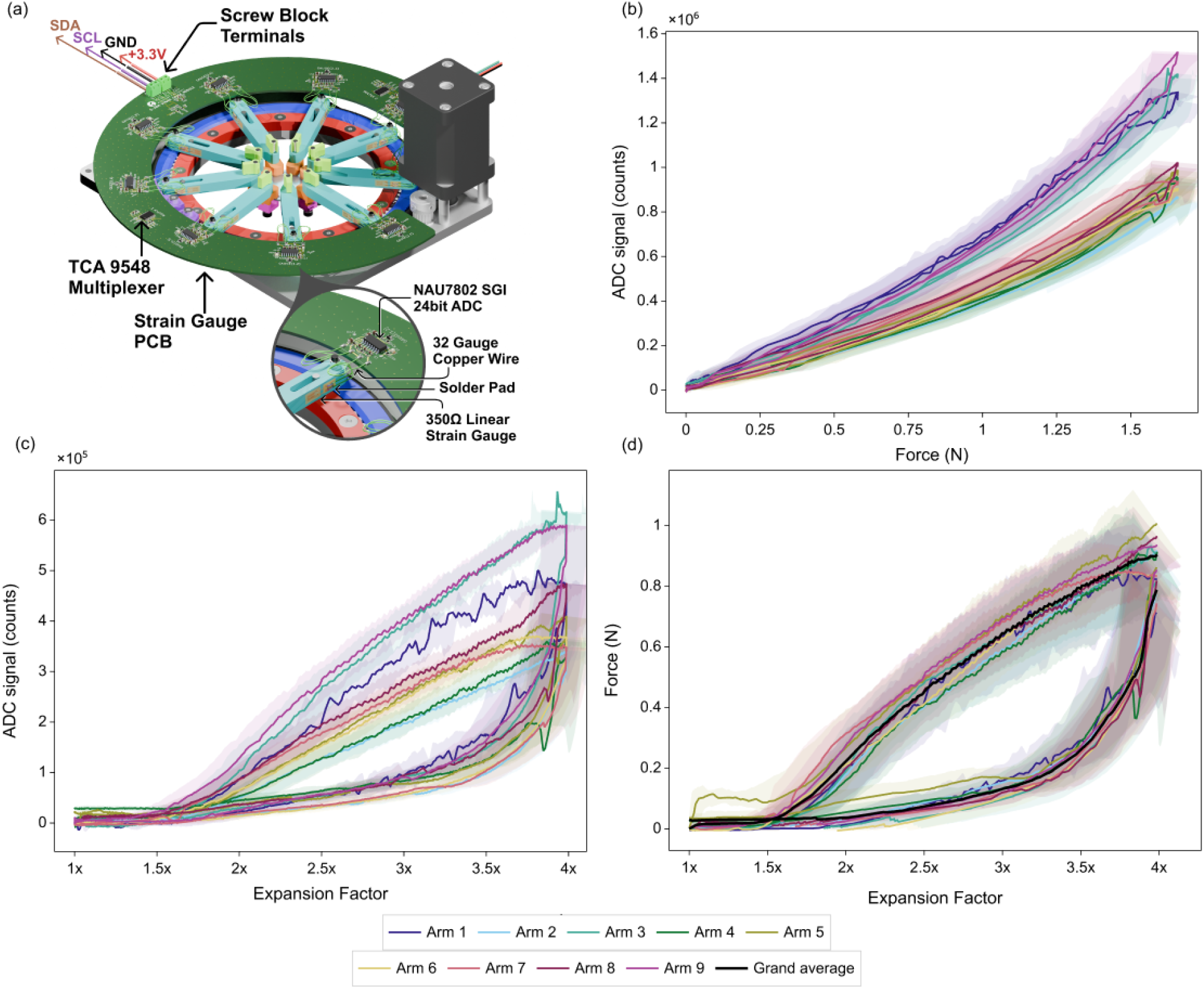
Strain-gauge force sensing in the TExM apparatus. (a) Load cells from linear gauges on 3D-printed arms, read out through a Wheatstone bridge and NAU7802 24-bit ADC, multiplexed over I^2^C by TCA9548 devices on the strain-gauge PCB; inset, a single gauge with solder pad and 32-gauge copper wiring. (b) Spring calibration: ADC signal versus applied force. (c) ADC signal versus expansion factor, per arm. (d) Force versus expansion factor for all nine arms, grand average in black; each curve is the mean of five trials. Shaded bands denote ±1σ. The curves form a closed hysteresis loop, loading above unloading across the mid-range and converging near maximum expansion, with the grand average reaching ∼0.88 N per arm at 4*x*. All arms agree within error, consistent with equibiaxial isotropy; residual deviations arise from angular tolerance, mounting variation, and sample preparation.

Each sweep expands the load from 1*x* to roughly 4.0*x*, held for a 5 second rest, then contracted back to 1*x*. Both expansion and hysteresis of the load were collected, characterizing the energy loss in the final force response curve. The ADC voltage was converted to force using a load of known force response and a spring of known constant, whose strain we measure directly in a secondary calibration measurement (Figure 5b). Interpolation between the calibration and the experimental hydrogel data included a moving window Savitzky-Golay filter [39], drift correction, and error propagation, as detailed in the SI section 3.2 and Figure S5.

#### 3.2.2 Isotropic Force Applied During Expansion

Incorporating strain gauge force sensors with the TExM stretcher enables real-time measurement of the force exerted on the hydrogel sample, allowing characterization of the polymer properties. The strain gauges were first calibrated, with Figure 5b showing the drift/offset-corrected ADC counts for two springs mounted on opposing arms, giving a consistent radial force on the arm of interest. The response is primarily monotonic and non-linear, with a mild superlinear tail, spanning 0 to 1.5 × 10^6^ counts over 0 to ∼ 1.6 N. A sharp dip appears at the intermediate rest period, with a lower but closely aligned contraction branch. Variations between arms arise from deviations in strain-gauge placement and infill structure, making each arm’s ADC scaling distinct but repeatable across trials, and thus is removed during interpolation.

Figure 5c shows the drift/offset-corrected ADC counts measured by the strain gauge under a hydrogel load and recorded over the expansion and contraction sweep of the TExM system. The ADC signal rises from near zero at 1× expansion to ∼ 3–6 × 10^5^ counts at 4×, with a slow initial regime and steeper accumulation beyond ∼ 1.75*x*. The maximum expansion used for the force measurement was limited to 4*x* due to the high sensitivity of the strain gauge, which produced high noise when the arms contacted one another at a maximum expansion of 4.2*x*. Using the spring calibration in Figure 5b, the ADC values for each arm with the hydrogel can then be converted to force.

Each individual arm applies a statistically identical force to the hydrogel. Figure 5d shows the force response for each arm averaged across five hydrogel trials, along with the grand average across all arms and trials. Similar to Figure 5c, the sweeps form a closed hysteresis loop: the expansion (loading) branch sits above the contraction (unloading) branch across the mid-range, with the two converging below maximum expansion (*E* ≈ 4.2*x*). The expansion branch follows a sublinear response, with the force climbing from ∼ 0.002 N at 1*x* to ∼ 0.86 ± 0.08 N at 3.75*x* and ∼ 1.01 N at *E* ≈ 4*x*, rising steeply from ∼ 1.75*x* onward before the rate of increase tapers off beyond ∼ 2.25*x*. The contraction branch follows a steep fall during the rest period and near full expansion, after which the force response follows a non-linear behavior, converging at the noise floor (≈ −0.0004 N to 0.033 N) for expansions below ∼ 1.5*x* . The grand average (black) tracks the per-arm curves closely on both branches, reaching a maximum of ∼ 0.9/0.1 N per arm at the set nominal maximum expansion of 4*x*, setting the representative per-arm loading at full expansion.

The large difference in the force response during expansion and contraction represents the viscoelastic properties of the hydrogel. The hysteresis is quantified by the enclosed loop in Figure 5d and the non-zero residual force on the contraction branch. Whereby the area encompassed by the two regimes is consistent with dissipative processes within the double-network hydrogel, including reversible ionic-network breakage and rearrangements. The initial plateau (Figure 5c, d) is due to the compression of the hydrogel during the mounting process. The compression from the grippers pushes material towards the center whereby the early portion of the expansion registers negligible tensile force until ∼ 1.75*x*, at which point the gel returns to its unstressed reference state. This is also reflected in the contraction curve which similarly meets this noise floor at the expansion interval (< 1.5*x*)

Despite the consistency in response across the arms and the grand average, several factors introduce the uncertainty into individual force measurements such as variation in strain gauge placement and the arm tolerance of the stretcher. Manufacturing processes such as machining and 3D-printing introduce the average arm angular deviation of 2°, as characterized previously (Figure 4), propagates directly into both the expansion and force axes. As outlined in the SI section 3.2.4 and Figure S6, this tolerance propagates the uncertainty in the expansion and, in turn, the force, giving the reported σ_*E*_ and σ_*F*_ on the final interpolated force-versus-expansion points shown in Figure 5. In addition, heterogeneity of the ionic crosslinking in the hydrogel network and irregular evaporation during sample handling results in variations in hydrogel stiffness. Moreover, the order and procedure of mounting of the hydrogel on the TExM system can influences the degree of contact and grip, resulting in heterogeneous stress distribution across the hydrogel before and during expansion. Despite these variations, the expansion and contraction sweeps of all nine arms remain statistically identical from one another, with residual deviations further minimized by averaging across multiple trials and supports the isotropy of the hydrogel expansion.

### 3.3 The TExM Iris Concentrates Surface Stress Near the Edge of Hydrogel with Uniform Stress at the Center of Hydrogel

#### 3.3.1 Finite Element Simulation of the TExM Stretcher

To understand the distribution of stress in our design, the TExM geometry and hydrogel substrate were modeled in COMSOL Multiphysics 6.2. The hydrogel mesh has 16 mm radius and 0.5 mm thickness with nine actuator cut-outs. A thickness of 0.5 mm was chosen since it improved the convergence of the simulation in relation to the actuator size and proportion while being close to the physical set-up. The cut-outs were arranged every 40° about the disk center at a radial center distance of 12.8655 mm, reproducing the nine-arm layout, with each actuator having a tangential width of 7.915 mm, a radial height of 4.557 mm, and an inner-corner fillet radius of 2 mm. These in-plane dimensions are shared with the two-dimensional model. Reaction forces were extracted by integrating over ball selections of radius 5 mm centered on each actuator.

The hydrogel was modeled as a compressible Neo-Hookean hyperelastic material with shear modulus *G*_g*el*_ = 1.724 kPa and bulk modulus *K*_g*el*_ = 16.67 kPa (Poisson ratio *v* = 0.45, nearly incompressible) and density 1000kg ⋅ m^−3^, with Lamé parameters *µ* = *G*_g*el*_ and 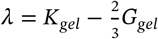 [40, 41, 42, 43]. Equibiaxial expansion was produced as prescribed radial displacements on the nine actuator side-wall faces, ramped linearly in time to a maximum of 30.88 mm (*u*_*x*_ = *dispmax, tcosθ*_*i*_, *u*_*y*_ = *dispmax, tsinθ*_*i*_, *u*_*z*_ = 0 with *θ*_*i*_ = *i* ⋅ 40°). Stiffness-proportional Rayleigh damping (*β*_*dK*_ = 0.015, *α*_*dM*_ = 0) was applied to support convergence. The problem was solved as a time-dependent study over *t* = 0 to 1 in steps of 0.05 (21 output points) with a relative tolerance of 1 × 10^−2^.

The finite element simulation reveals uniform, isotropic expansion near the center of the sample with stress concentrated by the grippers. Figure 6a shows the 2D expansion of the hydrogel to 4*x*, computed on a fine mesh under a plane-stress assumption with a fixed thickness of 0.5 mm. The von Mises stress is sharply localized to the edges of the hydrogel in the spans between neighboring grippers, because these regions carry the largest local strain as the material is drawn between adjacent clamps. The majority of the gel toward the center, where the specimen of interest sits, remains consistent in magnitude, indicating a uniform, isotropic expansion of the central region. The comparatively low and homogeneous interior field supports the assumption of equibiaxial isotropy used elsewhere in the analysis.

**FIGURE 6.**
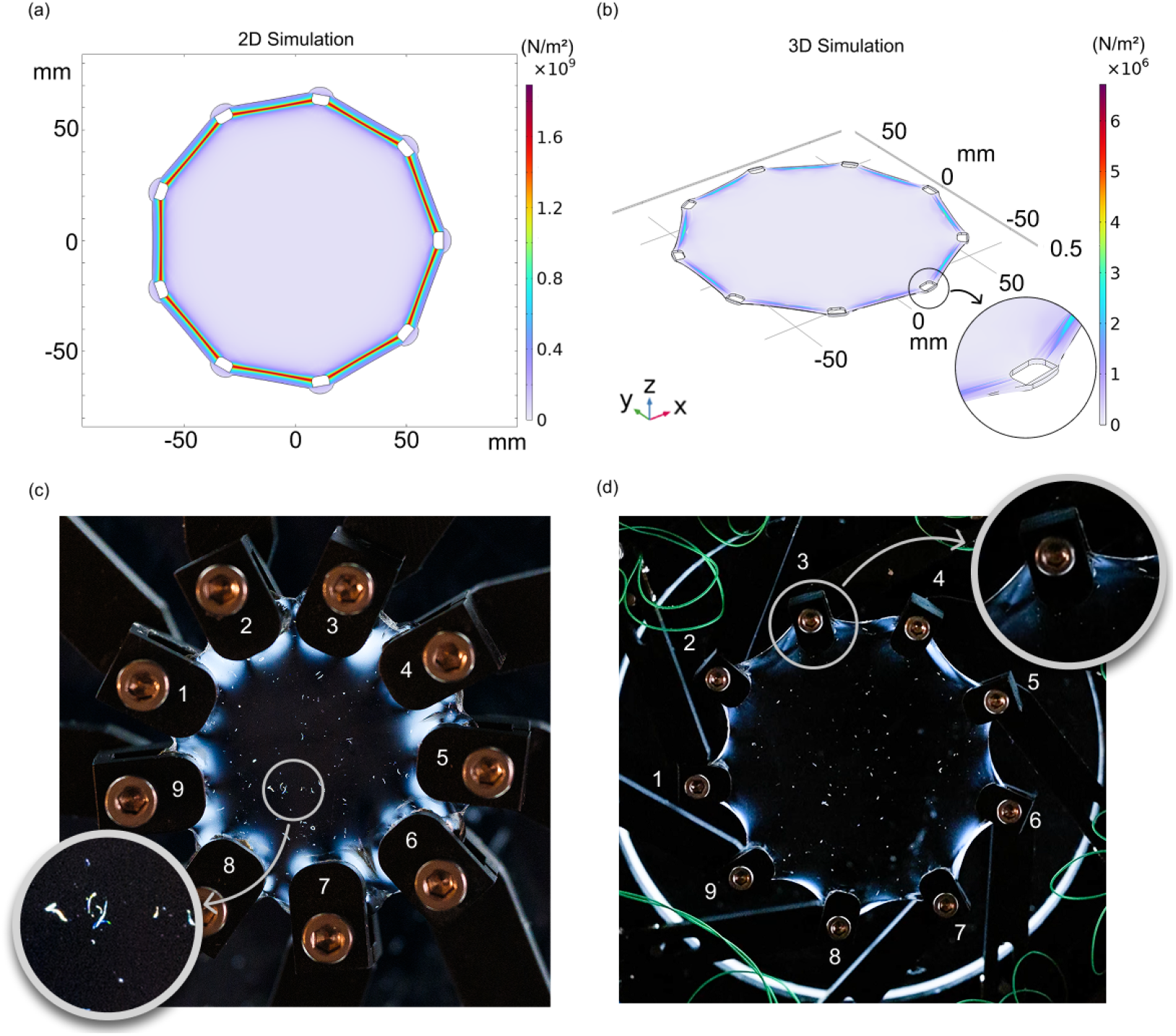
Finite-element and photoelastic characterization of stress in the hydrogel during expansion. (a) Two-dimensional surface and (b) three-dimensional volume plots of the von Mises stress computed in COMSOL for the 0.5 mm-thick gel held by nine radial grips, both at *t* = 1 s, the final frame of a *t* = 0–1 s simulated expansion corresponding to ∼ 4*x* linear expansion. Stress localizes along the free edges spanning adjacent grips while the interior remains comparatively uniform. (c, d) Plane-polariscope images of the gel in the nine-arm assembly (grips numbered 1–9) pre (c) and post-stretch (2.5*x*) (d). Inset (b): wrinkling at the gripper–gel interface. Inset (c): central pre-stretch region, showing only small birefringent debris and no fringe pattern. Inset (d): post-stretch gripper region, showing a bright birefringent band at the interface, consistent with the stress concentrations predicted in (a) and (b).

3D modeling which includes the hydrogel thickness produces similar qualitative results of stress being localized to the edge of the sample. Figure 6b shows the corresponding 3D expansion to 4*x*, computed on a fine three-dimensional mesh with the nine gripper portions removed and their boundaries displaced radially outwards at the same speed. The difference in the magnitude of the von Mises stress between the two plots suggests the 2D model constrains the gel to a uniform thickness whereby the applied load is carried entirely in-plane and concentrates at the gripper edges, driving the peak stress into the GPa range. However, in the 3D model the thickness is free to vary, allowing the hydrogel to thin and wrinkle out of plane near the grippers (inset Figure 6b); the load is then distributed throughout the thickness rather than along a single constrained plane, relieving these concentrations and lowering the peak stress by roughly three orders of magnitude, into the MPa range.

#### 3.3.2 Qualitative Experimental Evidence of Stress Distribution

To further support the finite element simulation results, a qualitative photoelasticity technique was used to visualize stress in the hydrogel during expansion through the process of stress-induced birefringence [44, 45]. The image was captured using a plane polariscope. A fixed polarizer sheet was mounted above an LED illumination pad to linearly polarize the transmitted light. A second polarizer (the analyzer), with an adjustable rotation angle, was fitted to a mirrorless camera mounted vertically on a tripod and aimed downward at the LED pad. The two polarizers were oriented with their transmission axes at 90° to one another (“crossed”), so that in the absence of any sample the analyses blocks essentially all the polarized light and appears dark (extinction). The TExM stretcher holding the hydrogel was positioned between the two filters, such that polarized light from the light box intercepts the gel before reaching the analyzer on the camera. Under load, the hydrogel exhibits stress-induced birefringence, where an otherwise optically isotropic (clear) medium becomes optically anisotropic in regions that are stressed. Any stress induced in the hydrogel alters the materials local polarizability with its own unique transmission axis and intensity related to the magnitude of the stress present. This introduces a relative phase retardation that re-orients the polarization state, which is observed by a greater transmission intensity in the analyzer. While the direct relationship between the magnitude of the stress induced to the polarizability of the hydrogel is unknown, this serves as a qualitative understanding to support the other techniques used.

Figure 6c shows the hydrogel in its contracted state at 1*x* expansion, with the nine gripper arms numbered for reference. Fine scratches visible across the surface of the gel indicate the localized miniature stress concentrations along the fractures (Figure 6c, inset). However, the scratches can potentially occur during preparation, handling and transfer of the hydrogel onto the stretcher. Large stress concentrations, indicated by the higher-intensity shades, are present around the boundary edges where the grippers meet the hydrogel. Heterogeneous stress concentrations stemming from the irregular clamping of the gel by the grippers which compresses the hydrogel unevenly and distributes it inconsistently across neighboring grippers are observed. The stress concentrations also vary from gripper to gripper, typically localizing around the geometry of each clamp: grippers 2 and 6, for example, show two to three distinct stress concentrations, whereas grippers 5 and 9 show more consistent concentrations around their filleted corners. These patterns push the hydrogel about the edges of the grippers, generating wrinkling between neighboring clamps, an effect qualitatively captured by the 3D simulation (Figure 6b), although the simulation does not reproduce the anisotropic stress distribution observed experimentally.

When expanded, stress concentration lies primarily along the boundary edges of the hydrogel and spans between neighboring grippers. Figure 6d similarly shows the stress distribution after expansion at approximately 2.5*x*. Moreover, the heterogeneity of the stress distributions still remains, exhibiting wrinkling in the stress field that is likely caused by minor folding of the hydrogel at the clamp and by imperfect mounting indicated on gripper 3. Such features alter the stress distribution and force response substantially from arm to arm, and will vary between trials as well.

Both simulation and experiment show that stress concentrates primarily toward the edges of the grippers and fades toward the center of the gel, where the least stress is observed. The photoelastic images do not, however, confirm the isotropy of the stress distribution at the center of the hydrogel, since the central field is not clearly visible. Nonetheless, the experimental images do support the trend predicted by the COMSOL simulations (Figures 6a, b), in which the highest von Mises stresses localize at the gripper boundaries while the center of the gel where the sample of interest would be located for imaging remains comparatively low and uniform. Differences are apparent between simulation and experiment, as the origin of the wrinkling in the simulation is a numerical artifact of mesh coarseness, whereas in the photoelastic images it results from gripper alignment and non-uniform traction on the gel.

### 3.4 Verification of Uniformity and Equibiaxial Stretch with Macroscale Fiducial Markers

To experimentally verify the equibiaxial and stretch uniformity generated by the TExM stretcher, macroscale rod-shape fiducial markers were drawn and video imaged on the hydrogel’s surface [16, 12]. Hydrogels were polymerized and soaked in calcium sulfate for 12 minutes, as described in our previous work [5], then the surfaces were patted dry. The gel was cut to an optimal diameter of 32 mm for sample loading. Fiducial markers were drawn via a 0.3 ultrafine sharpie marker due to the non-water soluble ink which prevents the ink from absorbing and dissipating into the hydrogel during expansion. To enhance consistency, a 3D-printed, 1 mm thick circular template with four rod-patterns, each at an equal distance of 2.5 mm from the center (rod dimensions = 3.5 mm × 1.5 mm) was used (Figure 7a). The rod dimension and spacing considered the marker head size and the area where cells are adhered to in the biophysical application of TExM [5]. Image acquisition was done using the same camera and settings as described in Section 3.1.

**FIGURE 7.**
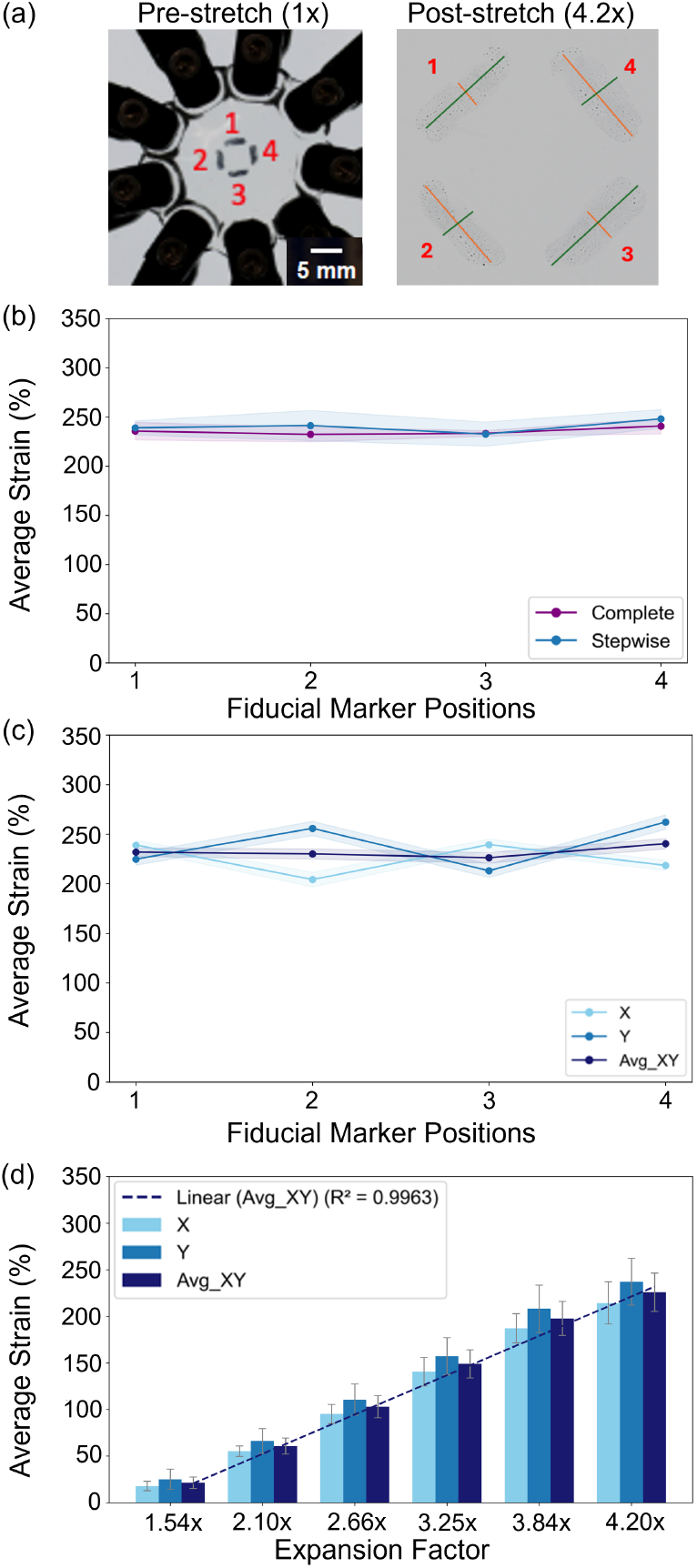
Validation of TExM stretcher illustrates uniform and equibiaxial stretch across (a) all four fiducial markers positioned on the center of the hydrogel. (b) There is no significant difference between two stretch conditions (complete and stepwise, n = 3 for each condition) while (c) shows strain% in X, Y, and average of both axes regardless of the stretch conditions (n = 10). The error bars represent standard error. (d) Change in strain % at each expansion factor during controlled stepwise stretch (n = 7) with speed of 0.1 cm/sec and 10 seconds dwell time after each stepwise expansion. Linear relationship between fiducial marker sizes and expansion factor is observed. There is no significant difference between stretching directions (x and y), thus indicating an equibiaxial stretch. Error bars represent standard deviation across seven gel samples (28 rod-shape markers in total)

Two conditions of expansion experiments, complete and stepwise stretch, were conducted to study the effect of stretching conditions on strain. The complete stretch condition is defined as a non-stop stretch from initial to maximum expansion factor (1*x* to 4.2*x*) at a speed of 0.1 cm/sec. Stepwise stretching, a condition used in cell expansion experiments [5] to progressive expansion at different factors, uses a similar speed but introduces a controlled stepwise expansion and a dwell time of 10 seconds between each step for hydrogel to come to equilibrium. Three sets of data for complete and seven sets of data of stepwise conditions were collected. The strain percentage was calculated from change in length of the markers along both X and Y axes relative to the grippers, following Equation 9. The axes are referenced to the movement of the grippers as the design introduced a maximum of ∼ 45 degree rotation of the arms which was transferred to the grippers. The change in length was measured manually by drawing line sections representing each axis on each rod (Figure 7a, post-stretch), using ImageJ software. Per each sampling, three line sections were drawn and averaged. A two-tailed student’s *t*-test of equal variance (unless otherwise noted) was used as a statistical method to validate equibiaxial stretch, stretch uniformity, and differences in stretch conditions from strain data. Statistical difference of the strains in x and y directions determine equibiaxial stretch while stretch uniformity is evaluated by strain values across each marker position on the hydrogel.

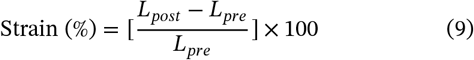

Complete or stepwise stretching does not have an effect in strain percent to the post-stretch samples (Figure 7b). A complete stretch would be used where only the maximum expansion is required to achieve super-resolution TExM, quickly going from 1*x* to 4.2*x* over 33 s, while the stepwise stretching is used to monitor cell response and any anisotropic expansion in real time during microscopy in smaller steps (0.01*x* to 0.5*x* over minutes). No significant difference is observed between these two modalities, with the *t*-test *p*-value = 0.26 (*p* > 0.05). Therefore, confirming the hypothesis that the stretching conditions do not affect resulting strain.

The expansion device applies ∼ 225% equibiaxial and uniform strain across the sample as assessed by macroscale markers. The average strain percent of each marker positions are not significantly different with *p* = 0.39 (Figure 7c). Thus, the fiducial rod markers experienced uniform strain applied by the stretcher.

Figure 7c also shows that the stretcher applies up to 240% average strain to the sample at maximum expansion (4.2*x*) by which the maximum strain for individual axes are 260% (y-axis) and 215% (x-axis), respectively.

The application of equibiaxial stretching to the sample throughout the expansion process can be determined by a continuous video of the full stepwise expansion. Figure 7d shows a linear relationship (*R*^2^ = 0.996) between average strain percent from both axes and expansion factor where strain% increases as the hydrogel is being expanded. The p-value from comparing average strain percent in X and Y axis at each expansion factor is 0.74, confirming no significant difference of strain in both axes. Thus, the TExM stretcher applies equibiaxial stretch and average strain up to 225% from both axes and maximum of 240% and 215% strain in Y and X axis, respectively.

Overall, section 3 verifies the performance of the TExM stretcher at the macroscale with average of 2° angular tolerance at the stretcher arms which apply consistent force to the hydrogel achieving uniform, equibiaxial stress and stretch at the center. With a calibrated, reproducible device, we now use the TExM stretcher for microscopic studies on an inverted fluorescence microscope. Microscope details can be found in SI section 4.

## 4 Automated Image Acquisition Software (AutoTracking)

An automated image acquisition software (“AutoTracking”) overcomes translational cell drift and focus shifts during hydrogel expansion, enabling cell tracking and eliminating manual focusing. This efficient workflow can preserve live-cell viability and shorten experiment time by reducing imaging duration. Figure 8a highlights the challenges that necessitate AutoTracking (analysis methods detailed in the SI section 5.1 and 5.2). Laterally, random shifts of X and Y positions of a feature over 100’s of pixels (10’s *µ*m) can occur during hydrogel expansion, as calculated by subtracting the centroid position of a feature relative to the image center (Figure 8a(i)). Axially, the stage z-position to collect an in-focus image is shown after each 0.04*x* expansion step (Figure 8a(ii); the increasing z-position over 100’s *µ*m confirms hydrogel thinning and underscores the necessity for custom autofocus software.

**FIGURE 8.**
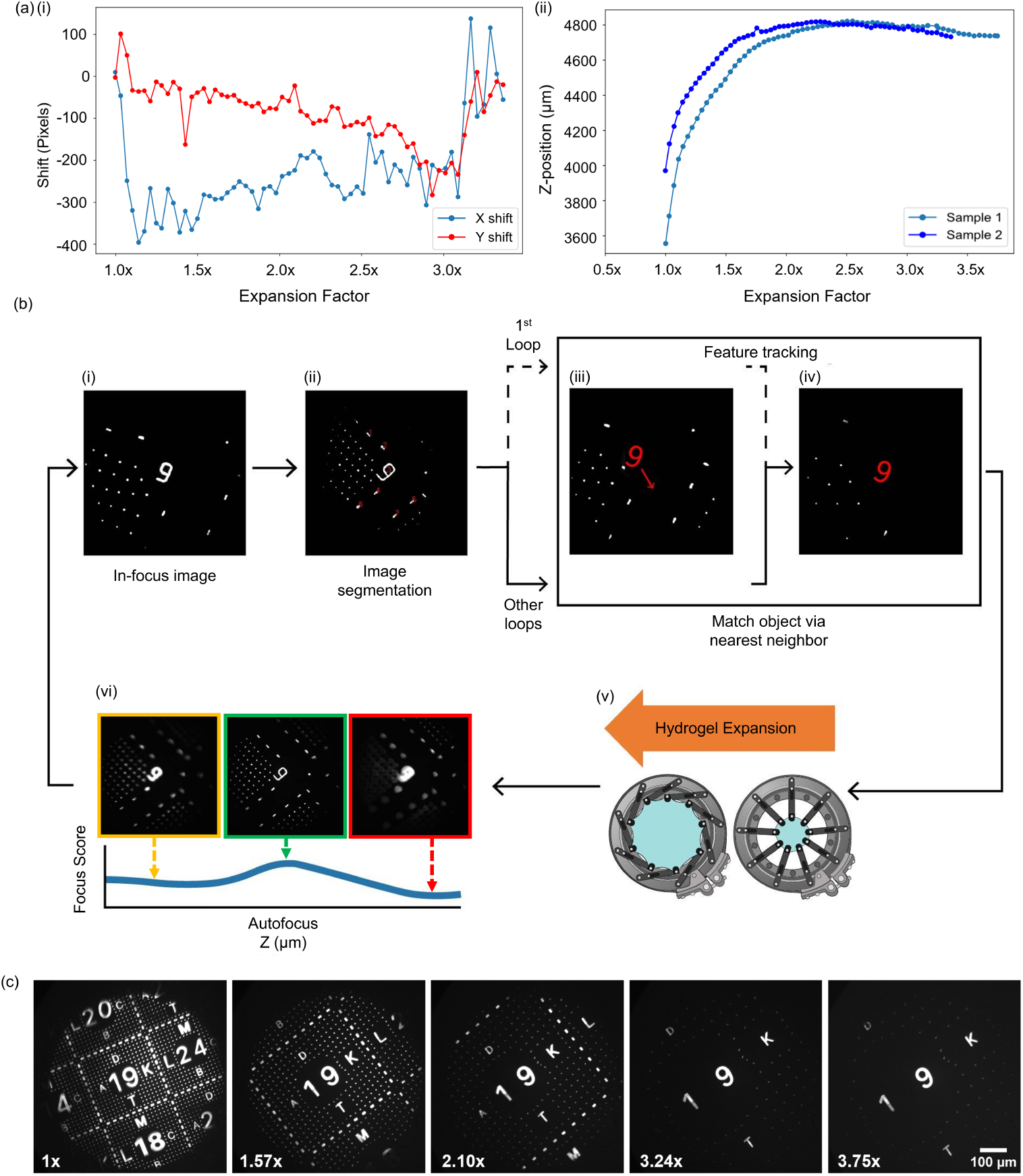
Real-time image acquisition using autofocus and tracking software. (a) Necessity of the software identified by (i) feature shifting along X and Y axes and (ii) changing of focal plane (z-axis) during expansion. (b) Workflow diagram of the software and (c) shows images of nanoscribe being tracked at each expansion factor

The AutoTracking software simultaneously controls the microscope’s piezoelectric stage, camera, and TExM stretcher while tracking fiducial markers, allowing for cell tracking throughout the expansion for final structural analysis of cells (Figures 8b, S8). The system integrates four core modules: microscope control, TExM stretcher control, autofocus, and feature tracking. Python serves as the central integration language, bridging the software languages of the Olympus IX83 microscope (Micro-Manager with a Java backend) and the TExM stretcher (C++ control script). Within the main script, the Pycro-Manager library handles microscope and camera operation [46], while pySerial manages serial communication with the stretcher [38]. The open source AutoTracking script is available at Github.

### 4.1 AutoTracking Software Description

The AutoTracking software starts with the user manually finding and selecting cells or features of interest with manual focusing (Figure 8b(i)) and input settings such as laser wavelength, exposure time, objective, and connection port number for the stretcher communication. The script then starts with first binarizing the in-focus image using adaptive thresholding of the average pixel intensity within the entire image plus one standard deviation. The regionprops Python library is used to extract five main characteristics of each feature (*f*): centroid, filled area, shape, Euler number, and perimeter [47]. Then, a size exclusion method is applied where an area threshold is set to only allow certain features to be annotated which remove noise (e.g., interference patterns, auto-fluorescence). The size-selected features are annotated with numbers (Figure 8b(ii)). The script then outputs an annotated image for the user to select and prompt one number associated with the interested feature to track throughout the expansion (Figure 8b(iii)). After the selection, the selected feature is centered in the field of view (Figure 8b(iv)), ready for the expansion.

Next, the script sends commands to the stretcher device to perform stretching at a set stepsize and speed (Figure 8b(v)). We found a small stepsize of 0.04*x* and speed of 0.01 cm/sec optimized effects of hydrogel dehydration, cell survival, and trackable shift of the interested features in X and Y planes. Slower stretching rate, smaller stepsizes, and slower speeds increase the accuracy of AutoTracking, however, it is more prone to hydrogel tearing and cell death in our non-environmentally controlled microscope setup prior to the full expansion. After each stepwise expansion, the hydrogel is set to rest for 10 seconds to reach equilibrium and ensure there is no movement to leverage the accuracy of the autofocus algorithm.

The autofocus algorithm is applied after each stepwise expansion to account for feature shift and hydrogel thinning. The Tenengrad algorithm, a Sobel gradient-based method known for accuracy and fast computational speed, is used in an OpenCV Sobel package [34]. A higher gradient value is present at the edges of features of in-focus images. The Sobel operator detects intensity variations along both horizontal and vertical directions then quantifies a focus score (*F*) by computing the mean gradient magnitude across all pixels within a kernel of 7 as shown in Equation 10 where *I* is image and *N* is the total number of pixels. We note that kernel size is an important parameter to adjust based on experiment settings and imaging conditions in order to achieve the most accurate in-focus image. The focus score, *F*, calculated over a range of axial steps of 3 *µ*m above and below the current z position with a region of interest (ROI) of 50% of the total lateral FOV at the center of the image to reduce processing time. Figure S9 shows an example of raw focus scores acquired from scanning through a z-stack images. The image that has the highest focus score is the in-focus image (Figure 8b(vi)). While acquiring and computing *F*, z-positions are saved and later send a command to move the stage to the z-position of the in-focus image via Pycro-Manager.

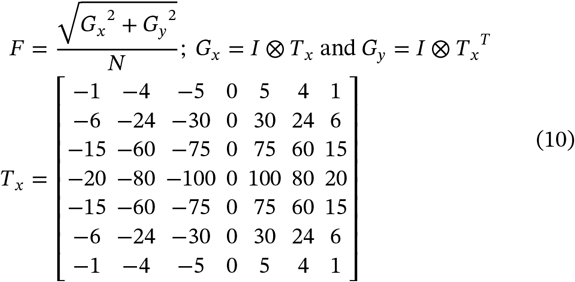

After acquiring the in-focus image from autofocus, feature tracking of the same selected-feature throughout the expansion relies on the nearest neighbor algorithm [48] of the previously described adaptive thresholding image segmentation. The five characteristics of the candidate features in the post-expanded image are compared with the saved characteristics of the selected feature (Equation 11 where *f* is five characteristics of each feature) by computing a user-defined weighted matching score (*w*). To prevent features with larger scales such as area from dominating the score, the absolute differences of all features are normalized using min-max normalization to a range of [0, 1] (Equation 12). We found that giving equal weight to all characteristics works in our case. Finally, the weight undergoes a dot product with normalized difference of each characteristic (Equation 13) to achieve a matching score for each candidate (Equation 14). The candidate feature that has the lowest score (the least difference in characteristics from target feature) is selected as the feature to track and the difference in distance from the center of the image and the centroid of the matched object is computed. Then, a command is sent via Pycro-Manager to move the microscope stage to ensure the feature of interest is centered before acquiring the final post-expansion image.

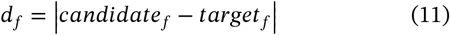

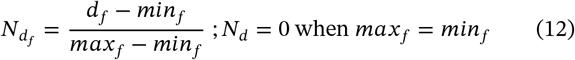

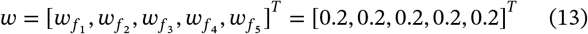

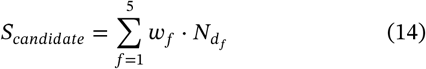

### 4.2 AutoTracking Software Verification with Microscopic Fiducial Markers

To verify that the software can successfully track the interested feature throughout expansion, a fluorescent fiducial marker made of Nanoscribe IP-Visio resin doped with Atto-633 fluorescence dye was used (fabrication method described in the SI Section 5.3 and our previous work [5]). Incorporating fiducial markers into the hydrogel improves AutoTracking accuracy due to sharp edges of the features compared to using less defined cell edges. The design of fiducial markers plays a crucial role in the success of tracking algorithms, where distinct features between adjacent characters significantly reduces decoding ambiguity and boosts tracking precision (Figure S10). The fiducial markers images were acquired using 633 nm laser at power of 2.5 mW and 50 ms exposure time using an Olympus IX83 microscope, which considered reducing the effect of photobleaching (Figure S11 describes imaging parameter optimization). The AutoTracking script acquires and tracks the interested feature while the stretcher applies a stepwise expansion of 0.04*x* with the speed of 0.01 cm/second which keeps the feature of interest in the FOV.

Successful AutoTracking image acquisition and feature tracking is shown from pre-stretch at 1*x* to post-stretch at 3.75*x* (Figure 8c). The fiducial marker of interest selected is the Nanoscribe patterned number 9. The AutoTracking algorithm is able to track the number 9 to the center of the FOV of every image across the full expansion (Figure 8c, left to right). Since the hydrogel needs to be fully taut to avoid any distortions of the fiducial markers, the pre-stretch expansion value started around the absolute position of the arms at 1.2*x* instead of 1*x*, which is then corrected for the post-expansion value in Figure 8c and limits full expansion to 3.75*x*. Yet, the algorithm would successfully work for a hydrogel mounted taut with the full 4.2*x* expansion range. Regardless, over 3.75*x* the “9” remains in the center of the FOV and is sharply in focus, demonstrating the success of the algorithm.

The combination of TExM stretcher and software allows for realtime tracking during expansion, in stark contrast to the before expansion, ex situ expansion, and after expansion imaging sequence required for osmotic ExM. Any distortion or structural changes of the sample during expansion can be identified with our real-time tracking capabilities (Figure S12).

## 5 Application: Fixed and Live Cellular Expansion

We further demonstrate an application of our TExM system to biological samples. NIH 3T3 fibroblasts were cultured on ITO coated cover glass within six well plates following standard cell culturing protocol and stained as described in the SI Section 6.1 and our previous work [5]. NIH 3T3 cells were fixed and embedded within the hydrogel, while live HeLa cells with endogenous fluorescent protein reporters were adhered on the surface of the hydrogel. Figure 9a shows pre and post expansion of fixed NIH 3T3 cells where single cell enlargement of ∼ 4*x* is observed along with details of the cellular structure on a wide field microscope. Detailed super-resolution analysis of the microtubular structures are provided in our previous work [5]. Multichannel images using different excitation wavelengths and labels allow visualization of both fiducial markers (used for AutoTracking) and the same cells throughout the expansion. HeLa cells were cultured and seeded onto the hydrogel according to the method discussed in the SI Section 6.2. Figure 9b shows pre and post expansion of live HeLa cells where robust cell-to-hydrogel adhesion is notably present before and after stretching. At 3.44*x* expansion cell cluster separation is observed potentially due to breakage of cell-to-cell adhesion, allowing visualization of cellular boundaries and enabling more accurate cell counting. Despite no significant increase in cellular area being observed (Figure S13), the precise, controllable capabilities of the TExM stretcher allows expansion to be stopped at any factors where cells are intact and higher resolution (fixed cells) or separation (live cells) images can be achieved. Thus, using TExM stretcher further promises future applications for dynamic cell culturing, mechanobiology study, expansion microscopy, and oncology (e.g., single-cell identification of cancerous cells or with drug modeling) [24, 49, 50, 51].

**FIGURE 9.**
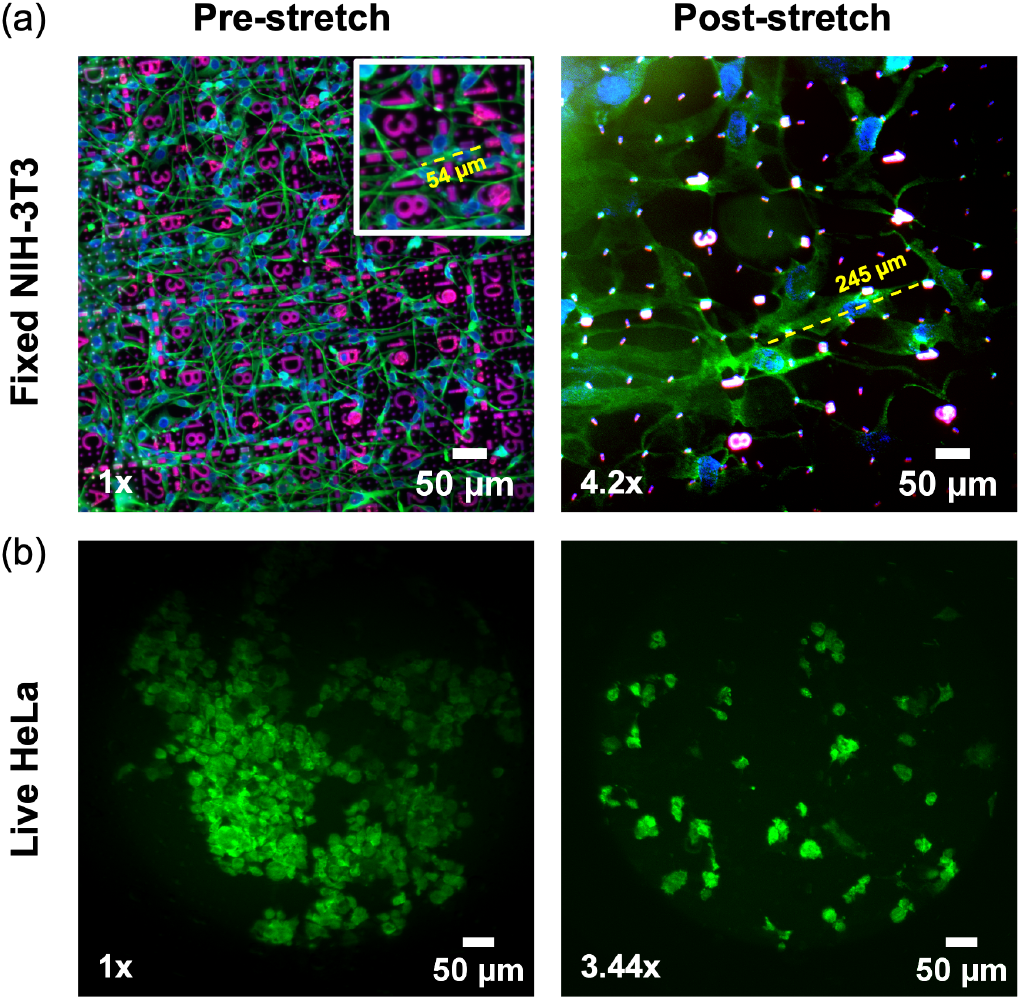
Pre and post-stretch images of (a) fixed NIH 3T3 fibroblast tagged with Alexa-488 for microtubule (green), Alexa-568 for nucleus (blue), and Atto-633 doped fiducial marker (red) and (b) live HeLa microtubule structure tagged with mClover3 fluorescence protein under 488nm laser (green). (a, inset) Zoomed-in region of the pre-stretch image shown in the post expansion image. The yellow line illustrates the length of the same single cell pre- and post-stretch which expands ∼ 4*x*. Note that the hydrogel needs to be taut for image acquisition, limiting the final expansion factor which is scaled accordingly in (b).

## 6 Conclusion

We designed and developed a novel, cost-effective, iris-based cell stretcher system for Tensile Expansion Microscopy (TExM) featuring integrated force sensors and real-time automated subcellular image acquisition with an AutoTracking software (Figures 2, 5, 8). Operating through a motor actuation mechanism controlled by a custom PCB, the device applies uniform equibiaxial stretch to a highly expandable double-network hydrogel substrate (Figures 6, 7). We validated the device on both fixed mouse fibroblast and live cervical cancer cell samples, demonstrating improved resolution and cellular expansion in fixed cells along with cell cluster separation in live cells (Figure 9). We believe our TExM system holds broad applications in super-resolution microscopy, mechanotransduction, dynamic cell studies, and polymer characterization. Furthermore, with slight modifications using arbitrary waveforms, the system could allow researchers to apply customized tensile force patterns while monitoring real-time effects on cellular structure through the AutoTracking software. The TExM system does have limitations, due to the multiple components which introduces operational complexity and optimization, requiring specific protocols for each part. Reoptimization may be needed when applying across different applications. Moreover, the substrate stiffness range is constrained by the maximum torque of the motor actuation system, preventing from stretching highly stiff hydrogels. To achieve higher expansion factors, a major redesign is required due to the limitations of the base plate size. Future directions including pursuing higher expansion factors (10*x*) comparable to the scales currently routinely achieved by osmotic ExM, application of TExM to diverse cell lines, developing compatibility with broader imaging platforms beyond Olympus fluorescence microscopes [52], and the integration of 3-in-1 directional control (uniaxial, biaxial, and equibiaxial) within a single platform. The TExM cell stretcher therefore can have an impact on understanding single cell mechanobiology and serve as in vitro model for organs that undergo tensile force such as skin, lung, and uterus.

## Author Contributions

R.A., V.V., D.L., M.Z., and L.K. contributed to conceptualization of the study and results interpretation. R.A. performed experiments and analysis for sections 3.1, 3.4, and 4. R.A. wrote sections 1, 2, 3.1, 3.4, 4, and 5. T.J. conducted experiments, analysis, and wrote sections 3.2 and 3.3. V.V. and D.L. performed fluorescence microscopy experiments and collected live and fixed cells images for section 5. V. Vakil collected the stretcher instructional assembly video. All authors reviewed the final manuscript.

## Acknowledgments

This research was supported by an Allen Distinguished Investigator Award, a Paul G. Allen Frontiers Group advised grant of Allen Family Philanthropies, a Scialog program sponsored jointly by Research Corporation for Science Advancement and the Gordon and Betty Moore Foundation and includes grant number 28410, and NIH NIGMS R35GM142466. We also thank Professor Yi Zhang and Dr. Dechen Fu for providing fluorescently labeled HeLa cells. The NIH 3T3 rd12 cells were a gift from Dr. Krzysztof Palczewski of UC Irvine. We also appreciate Cole Reinholt, Zhanda Chen, and Andrew Maytin for their early contributions to this project, Dr. Zechariah Pfaffenberger for consultation on the microscope, and Profs. Roberto Andresen Eguiluz and Divita Mathur for useful discussions and resources related to cell culture. Finally, we thank members of the entire Kisley lab and members of Prof. Laura Sanchez’ lab for discussion.

## Data Availability Statement

The data that support the findings of this study are available from the corresponding author upon reasonable request.

## Supporting Information

The Supporting Information section online provide stretcher design history, detailed iris stertcher data, hydrogel, cell culture, and microscopy methodology, derivation of error propagation for strain gauge force calculations, and additional data.

