## Supporting Information for "The Tensile Expansion Microscopy (TExM) cell stretcher: an iris expansion device integrated with automated real-time autofocus and tracking to super-resolve cells"

### 1 | Stretcher Design History

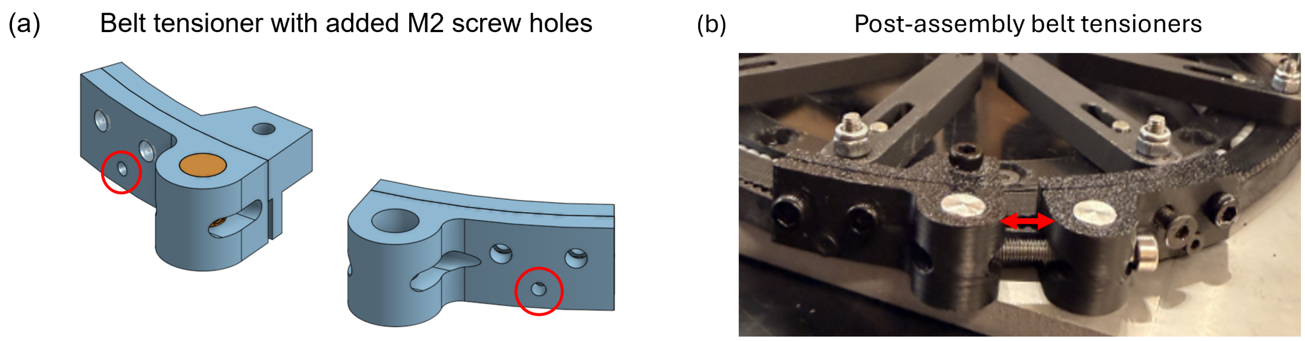


**FIGURE S1** | Design of belt tensioner mechanisms and post-assembly image. The belt tensioners undergo several modifications. (a) We found that having three screws instead of two to secure the belt in between the tensioners is crucial. Drilling a hole in the belt using a M2 screw (red circles)to fix it eliminates the issue of belt slacking. In addition, the tightness of the belt affects the torque transfer from stepper motor to the arms, the tighter the belt the more torque hydrogel receives to stretch them. Therefore, we adjust the belt to its maximum tightness. (b) Assembly of the belt tensioner on the TExM stretcher, it is important to leave a space between the two sides of the tensioners (red arrow) for belt tightness adjustment as the belt can be stretched or loosen over time.

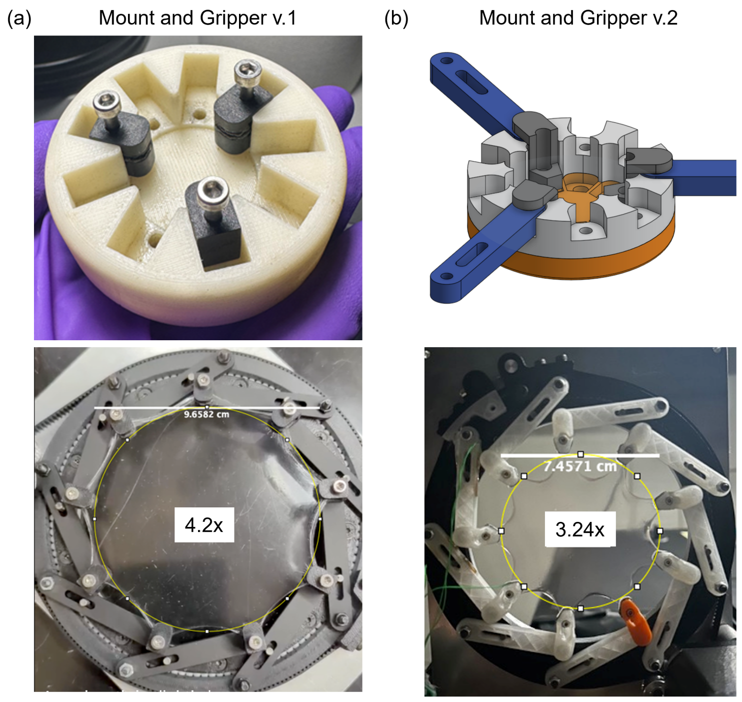


**FIGURE S2** | Different designs of gel mounting system and grippers were explored to determine their effects on maximum expansion factor while delivering the ease of sample preparation. The gripper version 1 (a) has more surface area to grip the hydrogel and shorter dimension compared to version 2 in (b), resulting in significant increase in expansion factor. However, mounting system of version 1 requires the user to lay and fix the hydrogel with nine screws in the mount first prior to lift up and transfer the hydrogel onto the stretcher device. Due to the sensitive nature of cellular samples cultured on the surface of the hydrogel, the lifting and transferring action poses high risk for damaging the sample leading to cells falling off of the surface and hard to handle for the user. The version 2 of the mounting system (b) involves a major redesign aiming for ease of use and eliminate the hydrogel lift up process. Thus, current stretcher design is built from those design ideas using the version 1 of gripper and version 2 of the mounting system to provide high expansion factor and maintain a user-friendly and creaful sample handling approach.

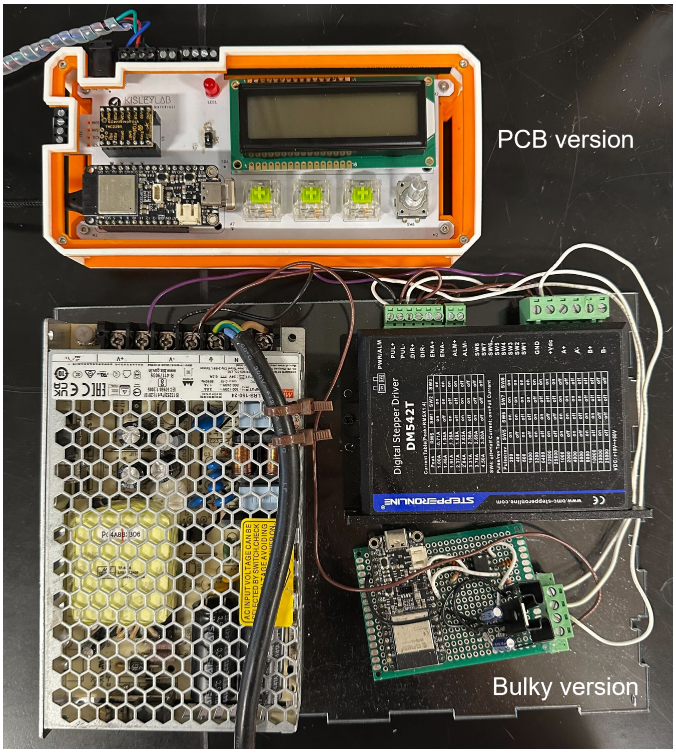


**FIGURE S3** | Different version of TExM control box. The first generation bulky control box (bottom) and PCB-based with LCD display (top). Another aspect contributing to a user-friendly design requirement is the design of the control box where compactness and modularity are the two crucial aspects. Our first generation control box shown below where it is heavy, bulky, not portable, and requires user expertise to set up. Indeed, we improve the design towards a modern PCB with controller integration.

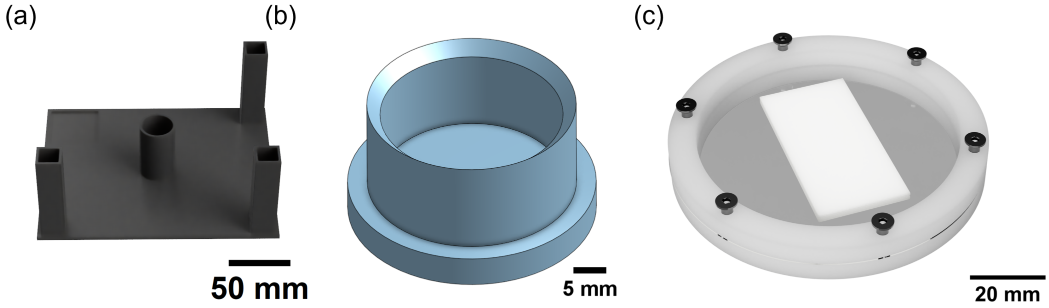


**FIGURE S4** | Other 3D printed components to support experiment set up (a) Stand used to support the stretcher while loading the hydrogel via the mounting system, (b) gel cutter used to cut the hydrogel to desired diameter, and (c) drying rings used to sandwich and dry cell on hydrogel samples after expansion for further imaging using 100$x$ oil objective.

### 2 | Hydrogel Preparation Method

The alginate-calcium/polyacrylamide double network (DN) hydrogel was used as a stretching substrate throughout the paper for fiducial marker imprinting and cellular adhesion. Hydrogel stock solutions were prepared at the following weight percent: Alginate (Alginic acid sodium salt from brown algae, CAS: 9005-38-3) at 2.91$wt.\%$, N,N'-methylenebisacrylamide (MBA; CAS: 110-26- 9) at 1.96$wt.\%$, Ammonium persulfate (APS; CAS: 7727-54-0) at 9.1$wt.\%$, and Acrylamide (AAm; CAS: 79-06-1) at 28.57$wt.\%$ (DN-Medium) and 50$wt.\%$ (DN-Stiff only used for fixed cells on hydrogel due to better transfer of cells, section 5 in main text). Due to the difference in AAm stock solution, the total polymer concentrations are 12.42% for DN-Medium and 23.36% for DN-Stiff.

To synthesize the hydrogel, Alg stock solution was measured and added to lass reagent bottle with screw cap (Fisherbrand™, Fisher Scientific). The Alg was dispensed from the stock solution using a 30 mL BD Luer-Lok™ plastic sterile syringe without a needle, with the weight precisely controlled on a digital scale. Then, AAm stock solution was added using a volumetric pipette. After that, the reagent bottle was sealed with a screw cap and wrapped in an aluminum sheet to prevent from light then subjected to magnetic stirring at a low velocity (50-100 rpm) for 45 minutes or until the solution is homogeneous. Then, APS stock solution was added using micropipettes then undergoes magnetic stirring for another 30 minutes to ensure homogeneity. Finally, the TEMED at $0.61 wt.\%$ (for DN-Medium) and $0.43 wt.\%$ (DN-Stiff) relative to the weight of AAm was added to the solution and subject to hand stirring using a strong magnet.

Once the precursor solution was homogeneous after TEMED addition, it was poured into the desired mold. A typical molding setup involved a circular silicone mold (50 mm in diameter and 1.5 mm in thickness) placed on a glass plate together with a substrate covered with a crystalline polyethylene terephthalate (PET) thin sheet (Amazon). The mold was then sealed by placing a second glass plate, also covered with a PET sheet, on top of the silicon mold, creating a "sandwich" structure which was critical to minimize the quenching of the free radical polymerization by oxygen in the air [1]. This assembly was then secured with large office binder clips. The precursor solution was placed in an oven at $35\,°C$ for 5 hours to achieve simultaneous free radical polymerization of AAm and covalent crosslinking with MBA, resulting in the formation of the PAAm covalently crosslinked network. The samples were then left to equilibrate overnight at room temperature to ensure complete network stabilization. After this period, the hydrogels were carefully peeled from the mold. With respect to specific application of the hydrogel, the protocol for soaking in a 20 mM calcium sulfate dihydrate ($CaSO_4 \cdot 2H_2O$, CAS: 10101-41-4, purity = 99% Thermo Fisher Scientific) differs accordingly. For DN-Medium hydrogel, after removing from the mold, the hydrogel was immersed in the calcium slurry for 6 minutes then carefully peeled off the substrate followed by another 6 minutes soaking. On the other hand, DN-Stiff hydrogel was peeled off the substrate first, then immersed in Calcium for 10 minutes, yielding the formation of the Alg-$Ca2^{+}$ ionically crosslinked physical network and thus Alg-$Ca^{2+}$/PAAm DN hydrogel. DN-Medium was used for section 3,4, and 5 (live cells) while DN-Stiff hydrogel was used only for section 5 fixed cells in the main text.

DN hydrogels composed of ionically crosslinked alginate and covalently crosslinked polyacrylamide represent a unique class of soft materials that combine exceptional extensibility with high toughness. Their outstanding mechanical performance originates from the complementary roles of the two interpenetrating networks. The brittle alginate network, formed through Calcium –mediated ionic crosslinks, acts as a sacrificial network that progressively dissipates mechanical energy through bond unzipping under deformation. In parallel, the highly stretchable covalently crosslinked polyacrylamide network maintains the structural integrity of the whole material by distributing stress and preventing catastrophic fracture. This synergy between the two interpenetrated networks enables the hydrogel to withstand large uniaxial and multi-axial deformations while remaining mechanically robust, making alginate–Calcium/polyacrylamide DN hydrogels an attractive platform for tensile expansion microscopy.

### 3 | Force Measurement using Strain Gauge

A strain gauge is a thin, patterned conductor (usually constant-an foil or sputtered metal film) bonded to an insulating backing. When the substrate below it stretches or compresses, the gauge's length and cross-section change, altering its electrical resistance. The fractional resistance change $\Delta R/R$ is proportional to the mechanical strain $\varepsilon$ by the gauge factor $GF$. A single gauge's resistance shift is tiny, often tens of micro-ohms for a full-scale load, so the device is wired into a Wheatstone bridge. To measure the small changes in voltages produced by the Wheatstone bridge configuration and derive some means of measuring the strain in the iris blade we need to use an analog to digital converter (ADC).

The gauges were mounted at the pin cavity, the point of maximum strain, to maximise the measurable response and improve sensitivity and ADC range, and the 32 AWG Gauge wires were soldered to jumper pads near the gauge itself for strain relief, reducing vibration transfer and the risk of joint breakage. The bridge is read by a NAU7802 24-bit ADC, which converts the small bridge voltages into a digital signal logged by an ESP32 microcontroller. The NAU7802 24 bit ADC contains four channels, allowing two bridges to be measured simultaneously. A custom PCB, laid out in KiCad and manufactured by JLCPCB, interfaces nine such ADCs with the ESP32 through two TCA9548A I2C multiplexers (routing four and five ADCs respectively); the board reduces the noise of loose jumper wires, arranges the nine chips into a compact ring around the iris arms, and gives a fixed, reproducible layout for consistent signal paths and grounding. The setup supports up to 18 strain gauges (two per blade across all nine blades) to measure both compression and tension on each blade. For this experiment, data collection was constrained to two strain gauges on a single blade. This configuration doubles the bridge's output voltage through the opposing resistance changes and cancels common-mode temperature drift for more reliable measurements.

### 3.1 | Iris Stretcher Data Collection

#### 3.1.1 | Assembly and Manufacture

To measure the small changes in voltages produced by the Wheatstone bridge configuration and derive some means of measuring the strain in the iris blade we need to use an analog to digital converter (ADC). The ADC acts as an interface to measure analog voltage signals as a digital signal which can be read and logged by the ESP-32 microcontroller. In this set-up I have used the NAU7802 24 bit ADC which contains four channels allowing for two bridges to be measured simultaneously.

A strain gauge is a thin, patterned conductor (usually constant-an foil or sputtered metal film) bonded to an insulating backing. When the substrate below it stretches or compresses, the gauge's length and cross-section change, altering its electrical resistance. The fractional resistance change $\Delta R/R$ is proportional to the mechanical strain $\varepsilon$ by the gauge factor $G_F$.

$$\frac{\Delta R}{R} = G_F\, \varepsilon \tag{1}$$

A single gauge's resistance shift is tiny, often tens of micro-ohms for a full-scale load, so the device is wired into a Wheatstone bridge. The gauges (DAOKAI BF350-3AA 350ohm High Precision Pressure Resistance Foil Strain Gauge, Amazon) were mounted at the pin cavity, the point of maximum strain, to maximise the measurable response and improve sensitivity and ADC range, and the 32 AWG Gauge wires were soldered to jumper pads near the gauge itself for strain relief, reducing vibration transfer and the risk of joint breakage. The bridge is read by a NAU7802 24-bit ADC (nuvoton) [2], which converts the small bridge voltages into a digital signal logged by an ESP32 micro-controller (Adafruit) [3]. A custom PCB, laid out in KiCad and manufactured by JLCPCB, interfaces nine such ADCs with the ESP32 through two TCA9548A I2C multiplexers (Texas Instruments) [4](routing four and five ADCs respectively); the board reduces the noise of loose jumper wires, arranges the nine chips into a compact ring around the iris arms, and gives a fixed, reproducible layout for consistent signal paths and grounding. The setup supports up to 18 strain gauges (two per blade across all nine blades) to measure both compression and tension on each blade. For this experiment, data collection was constrained to two strain gauges on a single blade. This configuration doubles the bridge's output voltage through the opposing resistance changes and cancels common-mode temperature drift for more reliable measurements.

**3.1.1.1 | Iris Stretcher Data Collection.** The stepper motor and strain-gauge readings are synchronized so that an averaged ADC voltage is recorded after each expansion step, linking every mechanical displacement of the iris blades to a corresponding strain measurement. Averaging multiple ADC samples per step reduces electrical noise and improves signal stability, while expanding the motor slowly preserves step definition and avoids vibration. The resulting ADC voltage versus step count dataset directly represents mechanical strain versus expansion, and forms the basis for the later Force versus Expansion analysis.

This is done through the Iris Stretcher Firmware which runs an experiment as a sequence of expansion targets, and during the run the experiment runner streams synchronized CSV rows over Serial.
Each row contains:
`exp` the experiment name
`t_ms` timestamp from `millis()` at sample acquisition
`steps` the motor's signed absolute step count
`target_ex` the current commanded expansion ratio
`state` M (moving), H (holding at a waypoint), or S (continuous strain stream)
`ADC1_mean`, `ADC1_std`, ..., `ADC9_mean`, `ADC9_std` mean and standard deviation of each ADC channel
At the beginning of each experiment the firmware initializes the 9 ADCs and tares their baseline reading. During an experiment run, for each step count, the strain array cycles through the nine NAU7802 ADCs, taking samples per chip, and reports the integer mean relative to the tare baseline along with its standard deviation calculated from the samples.

**3.1.1.2 | Python Serial Reader:.** A custom PyQt6 desktop application using PySerial [5] was developed as a counter part to the iris stretcher firmware to monitor and parse through the serial bus output of the microcontroller during experimentation and store the data collected. The application reads the USB serial CSV stream emitted by the ESP32, parses each row, and plots all nine ADC channels live with their mean and standard deviation against motor step count. Prior to data-collection the python script can be configured to three separate types of data collection as shown below:
**Hydrogel Expansion Dataset:** Raw ADC values vs step count while stretching the hydrogel.

**Spring Calibration Dataset:** Raw ADC values vs step count from all nine arms while stretching a spring of known spring constant.

**Drift/Offset Dataset:** Multiple raw ADC values vs step count while expanding the iris stretcher under no deliberate load. This captures both the drift of the ADC and the internal stresses that arise from the iris stretcher itself.

### 3.2 | Analysis pipeline

The goal of the pipeline is to convert the raw ADC counts of the strain-gauge measurements at a given hydrogel expansion into the force response curve, the process by which is not trivial. Since for each given measurement, the PCB reports the raw ADC counts from the NAU7802 SGI chip together with the motor step count we exploit the assumption that strain-gauge sensitivity and calibration to be common for every measurement made on the same setup. Hence, to recover the force response, we use a load of known force response, a spring of known constant, whose strain we measure directly in a secondary calibration measurement, and then interpolate the two datasets under a common strain.
This process can be broken into five stages which prepare and transform the hydrogel dataset to derive the force response for each given arm under a single trial as indicated by Figure S5.

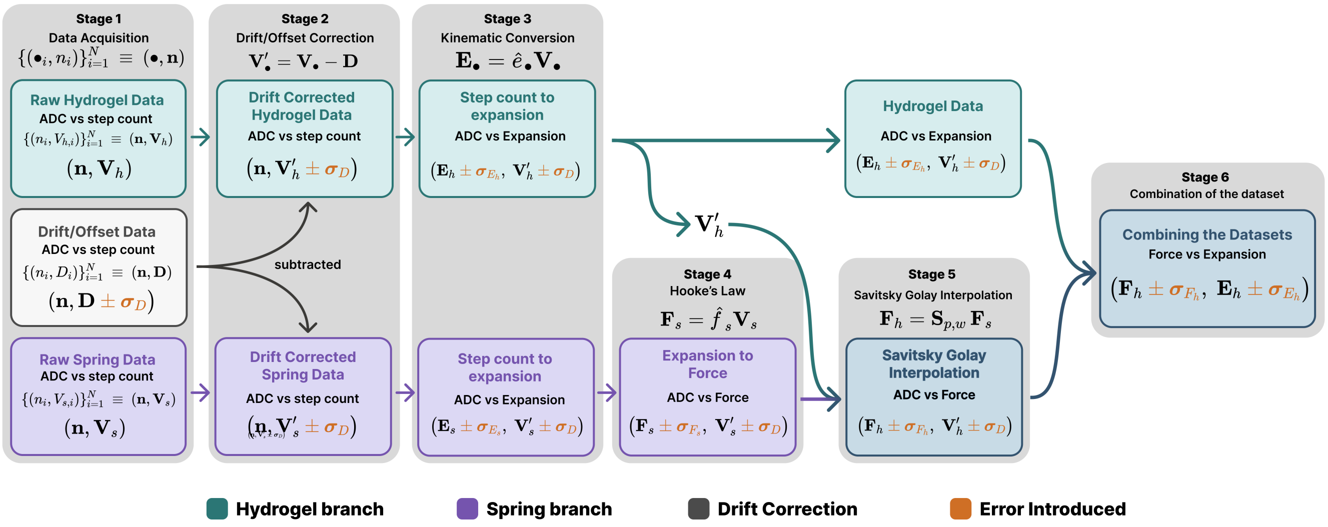


**FIGURE S5** | Diagram of the analysis pipeline for force measurement using strain gauge

#### 3.2.1 | Notation:

**Representing each per-arm Sweep:** To ensure consistency, all quantities below are given for a single representative *arm*. For a given sweep, all samples are indexed by $i = 1, \dots, N$, ordered by the commanded step count.
Each **dataset** is the full indexed family of samples, written as a set $\{(n_i, V_{h,i})\}_{i=1}^{N}$ or, equivalently, as aligned column vectors (as handled in Python). $\mathbf{V}_h$ denotes the per-sweep vectors, e.g. $\mathbf{V}_h = (V_{h,1}, \dots, V_{h,N})^\top$, while the corresponding individual element is represented as $V_{h,i}$. Relations between datasets are elementwise unless stated otherwise.
**Uncertainty convention.** To propagate the effective uncertainty of a derived quantity we use first-order (Gaussian) propagation. For $V_{h,i} = f(n_1, \dots, n_k)$ with independent inputs $n_i$, where $\sigma_{n_i}$ is the standard ($1\sigma$) uncertainty of $n_i$,

$$\sigma^2_{V_{h,i}} = \sum_i \left(\frac{\partial f}{\partial n_i}\right)^2 \sigma^2_{n_i}. \qquad (2)$$

Since the uncertainties are assumed small relative to the features of the dataset $V_{h,i}$, the first-order expansion in (2) is sufficient.

#### 3.2.2 | Stage 1: Data acquisition

Both tests use the same TExM system and record the strain-gauge response as a function of the commanded motor step count. A sweep produces $N$ samples; sample $i$ consists of the step count $n_i$ and the corresponding ADC reading $V_{\bullet,i}$ which

| Symbol | Meaning |
|---|---|
| $i = 1, \ldots, N$ | sample index within a sweep |
| $n_i$ ; $\mathbf{n}$ | motor step count (independent acquisition variable); its dataset |
| $V_{h,i}$, $V_{s,i}$ ; $\mathbf{V}_h, \mathbf{V}_s$ | ADC (strain-gauge) reading, hydrogel / spring; their datasets |
| $D_i$ ; $\mathbf{D}$ | drift (no-load) ADC reading; its dataset |
| $V'_{h,i}$, $V'_{s,i}$ | drift-corrected ADC reading, hydrogel / spring |
| $E_{h,i}$, $E_{s,i}$ | normalised expansion, hydrogel / spring |
| $F_{s,i}$ ; $F_{h,i}$ | spring force; hydrogel force (final, after interpolation) |
| $k$ ; $R_0$ | spring constant; geometric reference scale (Section, Phase 3) |
| $\Theta$ ; $\theta = \Theta - \frac{\pi}{2}$ | output-gear angle; mechanism angle |
| $r_0$, $r_p$, $r_{\text{pin}}$, $y_0$ | TExM system linkage lengths / offsets (Phase 2) |
| $c_0, c_1, c_2, c_3$ | linkage joint coordinates (Phase 2) |
| $E_0$ | relaxed (rest) reference scale for the expansion factor |
| $\mathcal{S}$ ; $\mathbf{H}$ | Savitzky–Golay operator; its smoother (hat) matrix |
| $w = 2m + 1$ ; $p$ | SG window length (half-width $m$); polynomial order |
| $W$ | SG window size, the quantity selected by SURE (Phase 3) |
| $\sigma_{\text{eff},i}$ | effective (both-axis) measurement uncertainty at sample $i$ |
| $\theta_D$ | per-arm angular manufacturing deviation |
| $\sigma_X$ | $1\sigma$ uncertainty of a quantity $X$ (e.g. $\sigma_{V_D}$, $\sigma_E$) |
| $\hat{(\cdot)}$ | a smoothed / fitted estimate |

for the interpolation will be the hydrogel test $V_{h,i}$ and the spring calibration test $V_{s,i}$. A third data set to account for the drift/offset $D_i$ which is outlined in following Section 3.2.3.
The three acquired datasets are therefore:

1. the **hydrogel test**, which loads the gauges through the stretched hydrogel,

$$\{(n_i,\, V_{h,i})\}_{i=1}^{N} \;\equiv\; (\mathbf{n}, \mathbf{V}_h);$$

2. the **spring calibration**, which replaces the hydrogel with a spring of known constant $k$ and provides the map that later turns an ADC reading into a force,

$$\{(n_i,\, V_{s,i})\}_{i=1}^{N} \;\equiv\; (\mathbf{n}, \mathbf{V}_s); \quad \text{and}$$

3. the **drift (offset) measurement**, the no-load baseline that is subtracted from the first two,

$$\{(n_i,\, D_i)\}_{i=1}^{N} \;\equiv\; (\mathbf{n}, \mathbf{D}).$$

#### 3.2.3 | Stage 2: Drift correction

A direct interpolation assumes that the ADC response for a given load depends purely on the load itself. Because the strain gauge is mounted directly on the arms of the apparatus, any load-independent deviation can be attributed to the stresses imposed by the TExM mechanism itself during expansion and contraction, rather than to the elasticity of the hydrogel. We therefore take a third measurement, which captures the drift and offset of the strain data during a no-load expansion, and subtract it from the other two denoted as $D_i$.
The drift dataset $\mathbf{D}$ is recorded with no load, capturing the baseline drift and offset of the gauges as the mechanism moves. Because it is acquired on the same step-count grid $\mathbf{n}$, it is subtracted sample-by-sample from each measurement stream to remove the baseline, giving the drift-corrected readings

$$V'_{h,i} = V_{h,i} - D_i, \qquad V'_{s,i} = V_{s,i} - D_i, \qquad \text{i.e.} \qquad \mathbf{V}'_h = \mathbf{V}_h - \mathbf{D}, \quad \mathbf{V}'_s = \mathbf{V}_s - \mathbf{D}. \tag{3}$$

Both corrected datasets remain in the form ADC versus step count; only the ADC axis has been shifted.
**Uncertainty introduced.** Subtraction (3) injects the uncertainty of the baseline into the corrected reading. Since we do not measure a define a given $\sigma_{V_h,i}$ & $\sigma_{V_s,i}$ we utilise the drift-induced ADC uncertainty $D_i$, derived from standard deviations of multiple trials, to define its error. Therefore the propogated uncertainty is:

$$\sigma_{V'_{h,i}} = \sigma_{V'_{s,i}} = \sigma_{V_D} \tag{4}$$

#### 3.2.4 | Stage 3: Step count to expansion

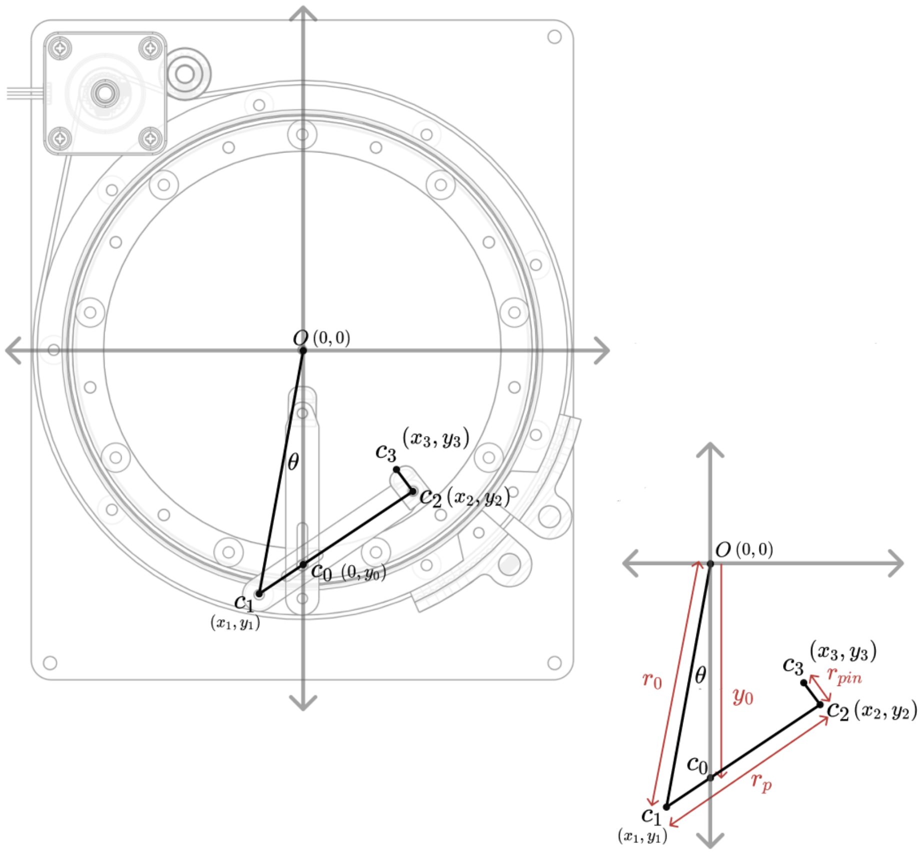


**FIGURE S6** | TExM system linkage geometry for the step-count-to-expansion mapping (Stage 3). The joints $c_0$–$c_3$, the driving radius $r_0$, arm length $r_p$, offset $y_0$, mechanism angle $\theta$, and gripper offset $r_{\text{pin}}$ are shown on the full mechanism (left) and the reduced linkage schematic (right).

Stage 3 maps the commanded step count $n_i$ to a normalised expansion $E_i$ through the kinematics of the TExM system linkage, leaving the (drift-corrected) ADC axis untouched. The result re-expresses each stream as ADC versus expansion: the hydrogel as $(\mathbf{E}_h, \mathbf{V}'_h)$ and the spring as $(\mathbf{E}_s, \mathbf{V}'_s)$. The derivation below is geometric and applies to either stream; the only difference between them is the reference scale $E_0$ used at the final step. This is shown in Figure S6.

**Step count to mechanism angle** The motor step count drives the output gear through a fixed transmission. With gear ratio $g$ and $p$ encoder pulses per revolution,

$$\Theta = \frac{2\pi n}{g p}, \qquad \theta = \Theta - \frac{\pi}{2}, \qquad s \equiv \sin\theta, \quad c \equiv \cos\theta, \tag{5}$$

where $\Theta$ is the output-gear angle and $\theta$ the mechanism angle measured from the horizontal.

**Linkage points** We place the linkage joints at

$$c_0 = (0, y_0), \qquad c_1 = (x_1, y_1), \qquad c_2 = (x_2, y_2), \qquad c_3 = (x_3, y_3), \tag{6}$$

with $c_1$ constrained to the driving circle of radius $r_0$,

$$x_1 = r_0 \cos\theta, \qquad y_1 = r_0 \sin\theta, \qquad x_1^2 + y_1^2 = r_0^2. \tag{7}$$

**Constraints** Two conditions fix $c_2$. First, the arm $c_1c_2$ is rigid, of constant length $r_p$:

$$r_p^2 = (x_1 - x_2)^2 + (y_1 - y_2)^2. \tag{8}$$

Second, $c_2$ lies on the straight line through $c_0$ and $c_1$, so

$$y_2 = \mu\, x_2 + y_0, \qquad \mu = \frac{y_1 - y_0}{x_1 - x_0} = \frac{y_1 - y_0}{x_1} \quad (x_0 = 0). \tag{9}$$

**Quadratic for** $x_2$ Substituting the collinearity relation (9) into the arm constraint (8) gives a quadratic in $x_2$,

$$r_p^2 = (x_1 - x_2)^2 + \big(y_1 - (\mu x_2 + y_0)\big)^2 \;\Rightarrow\; a\, x_2^2 + b\, x_2 + c = 0, \tag{10}$$

with coefficients

$$a = 1 + \mu^2, \qquad b = -2x_1 - 2y_1\mu + 2\mu y_0, \qquad c = x_1^2 + y_1^2 - 2y_1 y_0 + y_0^2 - r_p^2. \tag{11}$$

The physical (inner) solution is the minus root,

$$\Delta = b^2 - 4ac, \qquad x_2 = \frac{-b - \sqrt{\Delta}}{2a}, \qquad y_2 = \mu\, x_2 + y_0. \tag{12}$$

A pose with $\Delta < 0$ (or $c = 0$, an unreachable angle) is geometrically impossible and is returned as `NaN` and dropped from the curve.

**Expansion factor** The gripper radial position is the distance of $c_2$ from the centre,

$$\rho = \sqrt{x_2^2 + y_2^2} = \sqrt{a\, x_2^2 + 2\mu y_0 x_2 + y_0^2}, \tag{13}$$

the second form following from $y_2 = \mu x_2 + y_0$. Normalising by the relaxed (rest) reference scale gives the expansion factor,

$$E_x = \frac{\rho - r_{\text{pin}}}{E_0 - r_{\text{pin}}}, \qquad E_0 = r_0 - r_p, \qquad \xrightarrow{r_{\text{pin}}=0} \qquad E_x = \frac{\rho}{r_0 - r_p}. \tag{14}$$

By construction $E_x = 1$ at the rest pose. Evaluating (14) at each $\theta(n_i)$ produces the per-sample expansion; the streams become $(\mathbf{E}_h, \mathbf{V}_h')$ and $(\mathbf{E}_s, \mathbf{V}_s')$. The two streams use their own reference scales (the spring uses $E_0 = r_0 - r_p$, carried into Stage 4 as $R_0$).

The values for each of the parameters are listed below:

**TABLE 1** | Default TExM system-linkage geometry and drive parameters for the step-count-to-expansion mapping (Stage 3).

| Symbol | Description | Value | Unit |
|---|---|---|---|
| $r_0$ | driving radius | 7.25 | cm |
| $r_p$ | arm (pin) length | 5.32 | cm |
| $y_0$ | vertical offset of $c_0$ | -6.19 | cm |
| $r_{\text{pin}}$ | gripper pin offset | 0.766 | cm |
| $g$ | gear ratio | 12.7 | – |
| $p$ | pulses per rev | 1600 | – |

**Uncertainty introduced.** The arm-angle manufacturing deviation $\theta_D$ enters here as a *horizontal* (expansion-axis) uncertainty,

$$\sigma_E = \left|\frac{\partial E_x}{\partial \Theta}\right| \theta_D, \tag{15}$$

which collapses to zero as $E_x \to 1$. It is propagated in full in Section 3.1 of the main text.

#### 3.2.5 | Stage 4: Expansion to force

The spring stream is converted from expansion to force using the known spring constant; the hydrogel stream is carried forward unchanged. For a single arm the force is

$$F_s = 4\cos^2\!\Big(\frac{\pi}{18}\Big)\,(E_s - 1)\, R_0\, k, \tag{16}$$

where $k$ is the spring constant, $R_0 = E_0^{(3)} = r_0 - r_p = 0.0193\,\text{m}$ is the Stage-3 reference scale (with $r_{\text{pin}} = 0$), and $4\cos^2(\pi/18)$ is the fixed geometric factor of the nine-bladed TExM system. The factor $(E_s - 1)$ makes $F_s = 0$ at rest. These are shown in Figure S7.

Equation (16) turns the spring stream into ADC versus force, $(\mathbf{F}_s, \mathbf{V}'_s)$; equivalently it defines the per-arm calibration that maps an ADC reading to a force. The hydrogel stream remains $(\mathbf{E}_h, \mathbf{V}'_h)$. Both streams now share the ADC reading as a common variable, but in different pairings:

$$\text{hydrogel: } (\mathbf{E}_h, \mathbf{V}'_h), \qquad \text{spring: } (\mathbf{F}_s, \mathbf{V}'_s). \tag{17}$$

The spring constant (94135K328, McMaster-Carr) is defined in table 2.

**TABLE 2** | Spring calibration constant used to convert the spring stream to force (Stage 4).

| Symbol | Description | Value | Unit |
|---|---|---|---|
| $k$ | spring constant | 12.259 | N/m |

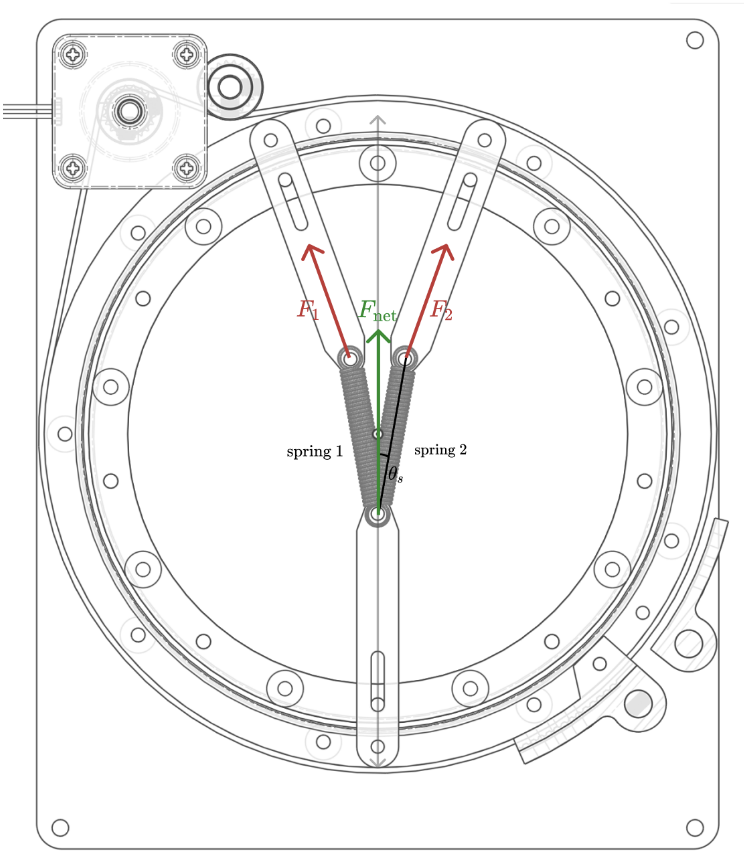


**FIGURE S7** | Spring calibration configuration on the TExM system, with two springs of known constant $k$ mounted in place of the hydrogel across a pair of adjacent arms. Each spring pulls along its arm with force $F_1$ and $F_2$; their components along the central axis sum to the net calibration force $F_{\text{net}}$, while the transverse components cancel by symmetry. The half-angle between each spring and the central axis is $\theta_s = \frac{\pi}{9}$, which sets the geometric projection factor in the spring-to-force conversion of Stage 4.

#### 3.2.6 | Stage 5: Savitzky–Golay interpolation onto a common ADC axis

**Utilising a common strain-gauge calibration.** The conversion is possible only because both sweeps are read out by the *same* strain gauge, with the same sensitivity and calibration. The spring calibration therefore does more than record data: it measures, for this exact gauge, the force that produces any given drift-corrected ADC reading. Put differently, the spring pairs $\{(V'_{s,i}, F_{s,i})\}$ trace out a single force-versus-ADC map

$$F = \varphi(V'), \qquad \varphi : \text{ADC reading} \longmapsto \text{force}, \tag{18}$$

which is a property of the gauge, not of the spring. The hydrogel sweep, read by that same gauge, must obey the same map: whatever load the hydrogel applies, an ADC reading of $V'_{h,i}$ corresponds to a force $\varphi(V'_{h,i})$. Converting the spring force vector $\mathbf{F}_s$ into the hydrogel force vector $\mathbf{F}_h$ is thus nothing more than evaluating this shared map at the hydrogel's ADC values. The only concern is the two sweeps do not sample the same $V'$, so $\varphi$ must be reconstructed from the discrete spring pairs and evaluated off-grid. That reconstruction is the interpolation, for which we use a Savitzky–Golay fit.

**The Savitzky–Golay operator.** A Savitzky–Golay (SG) filter fits a low-order polynomial to a moving window of points and reads off the fitted value at the query point [6]. It is a *linear* operator, so applying it to a dataset is a matrix–vector product. Writing the spring force on its own ADC grid as $\mathbf{F}_s$, the smoothed/resampled force is

$$\hat{\mathbf{F}} = \mathbf{H}\,\mathbf{F}_s, \qquad \hat{F}_i = \sum_j H_{ij}\,F_{s,j}, \tag{19}$$

where the smoother (hat) matrix $\mathbf{H}$ follows from the (weighted) local least-squares fit as defined by the

$$\mathbf{H} = \mathbf{A}\big(\mathbf{A}^\top\mathbf{W}\mathbf{A}\big)^{-1}\mathbf{A}^\top\mathbf{W}, \qquad \mathbf{W} = \operatorname{diag}\!\big(\sigma_{\mathrm{eff},i}^{-2}\big), \qquad \operatorname{tr}(\mathbf{H}) = \sum_i H_{ii}. \tag{20}$$

Here $\mathbf{A}$ is the Vandermonde design matrix of the local polynomial basis (order $p$), and the window length $w = 2m+1$ sets the bandwidth of $\mathbf{H}$ (a wider window is a wider band) [6]. Resampling the spring calibration at the hydrogel ADC values $\mathbf{V}'_h$ gives a force at every hydrogel sample,

$$F_{h,i} = \hat{s}\big(V'_{h,i}\big) = \big(\mathbf{S}\,\mathbf{F}_s\big)_i, \qquad \mathbf{S} = \mathbf{S}(\mathbf{V}'_s, \mathbf{V}'_h) \in \mathbb{R}^{N_h\times N_s}, \tag{21}$$

with $\mathbf{S}$ the (rectangular) SG resampling operator built from the spring grid and evaluated on the hydrogel grid. Equation (21) is the explicit $\mathbf{F}_s \to \mathbf{F}_h$ conversion: $\hat{s}$ is the smoothed, data-driven estimate of the calibration map $\varphi$ of (18), and applying it at each hydrogel ADC reading transfers the spring's known force onto the hydrogel samples. Nothing about the hydrogel's material enters here; the spring supplies the force scale and the hydrogel supplies only the ADC values at which that scale is read off.

**Window selection (SURE).** The window size $W$ is chosen objectively by minimising Stein's Unbiased Risk Estimate [7], which estimates the deviation of the fit from the true signal using only the measured error,

$$\mathrm{SURE}(W) = \underbrace{\sum_i \big(F_i - \hat{F}_i\big)^2}_{\text{fit to data}} - \sum_i \sigma_{\mathrm{eff},i}^2 + \underbrace{2\sum_i \sigma_{\mathrm{eff},i}^2\,H_{ii}}_{\text{complexity penalty}}. \tag{22}$$

The leverage $H_{ii}$ measures how much each point drives its own fitted value, so $\operatorname{tr}(\mathbf{H}) = \sum_i H_{ii}$ is the effective number of degrees of freedom: a narrow window gives high leverage ($\operatorname{tr}\mathbf{H} \to N$) and a large penalty, a wide window spreads the influence ($\operatorname{tr}\mathbf{H} \to p+1$). The correction is exact rather than heuristic, since $\sigma_{\mathrm{eff},i}^2 H_{ii} = \operatorname{Cov}(\hat{F}_i, F_i)$ and Stein's identity sets the total optimism to $\sum_i \operatorname{Cov}(\hat{F}_i, F_i)$. In practice $\mathbf{H}$, and hence $t = \operatorname{tr}(\mathbf{H})$, is read from the `interp_savgol` routine, and $W$ is taken at the minimiser of (22) [7].

The effective variance combines the uncertainties on both axes (the Orear effective-variance construction),

$$\sigma_{\mathrm{eff},i}^2 = \sigma_{F,i}^2 + \left(\frac{\partial F}{\partial V}\right)^2 \sigma_{V,i}^2. \tag{23}$$

**Result: pairing force with expansion.** The interpolation has now given every hydrogel sample a force $F_{h,i}$, while Stage 3 already gave that same sample an expansion $E_{h,i}$. Both quantities carry the same index $i$ and both descend from the one hydrogel ADC reading $V'_{h,i}$: the expansion through the kinematics of Stage 3, and the force through the calibration map of (21). They can therefore be paired sample-by-sample, which eliminates the ADC variable and leaves force as a function of expansion,

$$V'_{h,i} \xrightarrow[\text{Stage 3}]{\text{kinematics}} E_{h,i}, \qquad V'_{h,i} \xrightarrow[(21)]{F_{h,i}=\hat{s}(V'_{h,i})} F_{h,i} \qquad \Rightarrow \qquad (E_{h,i},\, F_{h,i}). \tag{24}$$

Collecting over all samples yields the final per-arm curve,

$$\boxed{(\mathbf{E}_h,\ \mathbf{F}_h)} \qquad \text{(force versus expansion).} \tag{25}$$

The drift-corrected ADC reading $V'_{h,i}$ has served only as the intermediate variable that links expansion to force: the kinematics convert step count to $E_{h,i}$, the spring calibration converts the ADC reading to $F_{h,i}$, and pairing the two at matched $V'_{h,i}$ removes it from the final result.

### 4 | Fluorescence Microscopy

Imaging was performed on an inverted fluorescence wide field microscope [8] with additional lasers and brightfield module. The home-built super-resolution fluorescence microscopy setup used Olympus IX-83 microscope body, brightfield light pillar with halogen lamp (IX3-ILL, U-LH100-3-7), and free-standing optics for excitation by either a 100 mW 488 nm (Toptica iBeam Smart), 100 mW 561 nm (CNI), or 100 mW 633 nm (Toptica iBeam Smart) laser The laser power was controlled using Toptica software. The beam passed through optical path then reached dichroic mirror (either Chroma ZET488rdc, ZET561rdc, or ZET633rdc for 488, 561, or 633 nm excitation, respectively) which directed the beam to a 20x/0.5 NA air objective(Olympus UPLFLN). The laser power densities at the stage are 2.5 $mW/cm^2$ for AutoTracking fiducial markers and 5 $mW/cm^2$ for cells images. The sample position was controlled by xy piezostage (Olympus IX3-SSU) and objective positioner (Olympus IX3-D6RES). 16-bit images were acquired through a Photometrics Prime 95B 22 mm scientific CMOS camera after passing through additional emission filters (Chroma ZET488NF, ZET561NF, ZET635NF for 488, 561, or 633 nm excitation, respectively) at 50 ms exposure time for AutoTracking fiducial markers and 150 ms for all cells-related images.

### 5 | Auto-tracking software

#### 5.1 | Analysis Method for Feature Shifting during Expansion

To better quantify and visualize the effect of feature shifting during expansion, following each stepwise stretching (step size = $0.04x$), a post-stretch image containing a non-centered interested feature was collected to extract the centroid of the interested feature. After collecting all post-stretch images until complete expansion, all images were imported into a Python script for analysis. The Python script used an image segmentation algorithm similar to the AutoTracking software to annotate and extract feature characteristics in the image then the user manually selected the interested feature. The regionprop library then outputs a centroid (X and Y positions relative to the image pixel) of the selected feature. Then, the shifted pixel was obtained by subtraction between the centroid of the selected features and center of the image. The process was repeated for each post-stretch image throughout the full expansion. Figure 8a(i) in main text presents a plot of shifted pixels at each stepwise expansion.

#### 5.2 | Analysis Method for Focus Drift during Expansion

Focus drift in z position was determined by collecting microscope stage metadata of in-focus image during expansion image acquisition. As the stretcher expands hydrogel in a step of $0.04x$ equibiaxially, autofocus was applied to obtain in-focus image post-stretch. AutoTracking algorithm collected Metadata associated with stage positions. The raw z-position data of each post-stretch image were then plotted against expansion factor as shown in Figure 8a(ii) in main text.

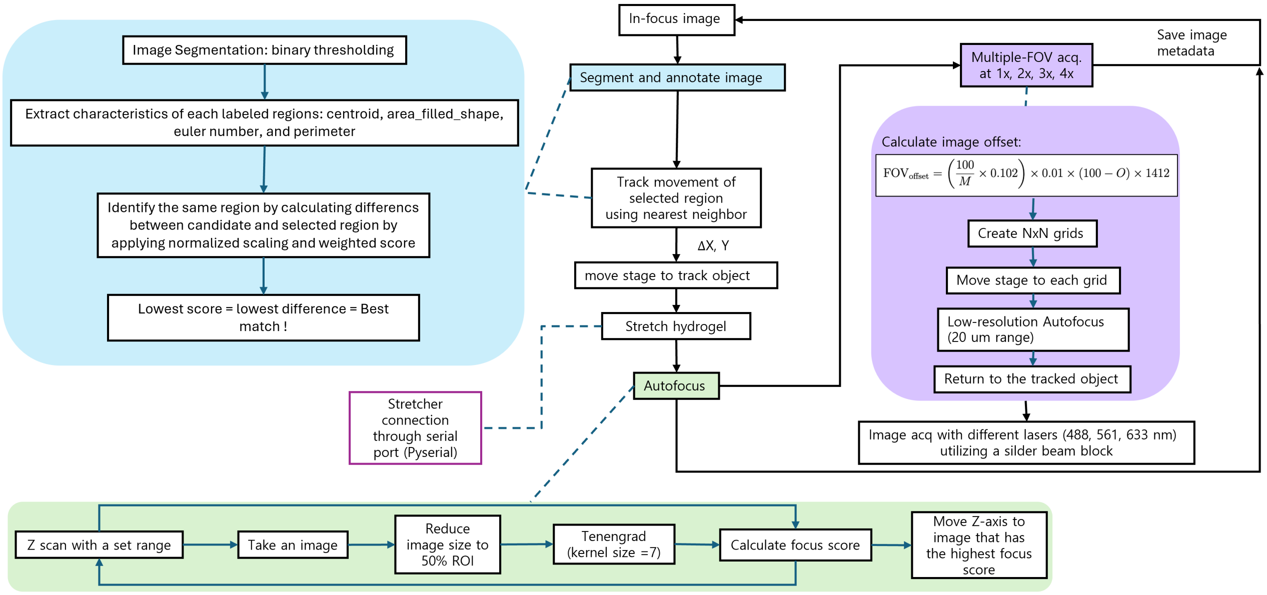


**FIGURE S8** | Detailed workflow diagram of the AutoTracking software with the additional function for multiple field of view acquisition for interested regions.

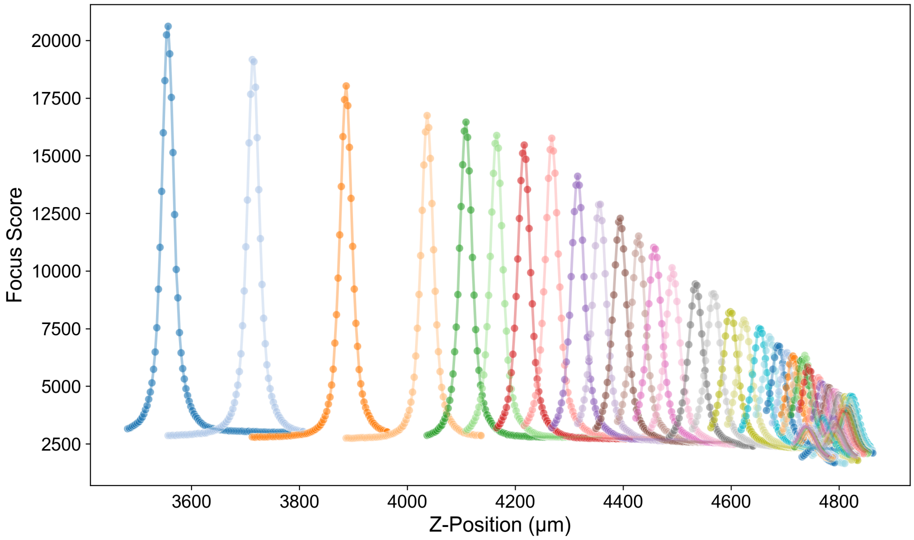


**FIGURE S9** | Raw focus score obtained applying autofocus to a range of z-scan images. Different colors showing each set of z-scan after each stepwise expansion to find in-focus images. The peak of focus score in each set indicate the z-position of the in-focus image. As the sample expands, the number of fiducial markers decrease per one field of view, meaning there are less sharp edges in an image. Thus, there is a decreasing trend of the focus score across the expansion.

### 5.3 | Fabrication and transfer of fiducial markers onto hydrogel

Fiducial markers were used to verify AutoTracking software. Fabrication and transfer steps are based on the protocols established in the TExM paper[9] and modified for specific applications. The micron-size markers were printed following a two-photon lithography technique on clean silicon wafer substrate (for live cells) or an ITO coated 1.5 glass coverslip (for fixed cells) using the Nanoscribe Photonic Professional GT2 machine. IP-Visio resin, a low auto-fluorescence negative tone photoresist, doped with 0.1 mM Atto-633 (data without cells) or 0.1 mM Abberior Star Red (data with cells) was chosen to print the fiducial markers. Dip-in laser lithography mode was used to print the pattern on a wafer with optimized printing parameters (power scaling = 1, laser power (%)= 80, scan speed ($\mu m/s$) = 2000, and Galvo Acceleration ($V/ms^2$)=0.4). The features were developed in propylene glycol methyl ether acetate (PGMEA; CAS:108-65-6, Sigma Aldrich) for 20 minutes and cleaned in isopropyl alcohol (IPA; CAS: 67-63-0, Sigma Aldrich) for 5 minutes. The patterned substrates were then dried using a nitrogen gun and stored in the dark until ready to use.

The fiducial markers on the substrate were ready for transfer onto the hydrogel. DN-Medium hydrogel on Silicon wafer was used for section 4 and 5 (live cells) while DN-Stiff hydrogel on ITO coverslip was used for section 5 fixed cells in the main text. The hydrogel synthesis is described in described in 2. After having fiducial markers on hydrogel, the hydrogel was loaded onto the stretcher device according to the protocol previously mentioned in Section 2.1 in the main text then placed onto the microscope stage. After connecting to power and establishing connectivity with the PC, the Micromanager software should be left running in the background while the AutoTracking script is running.

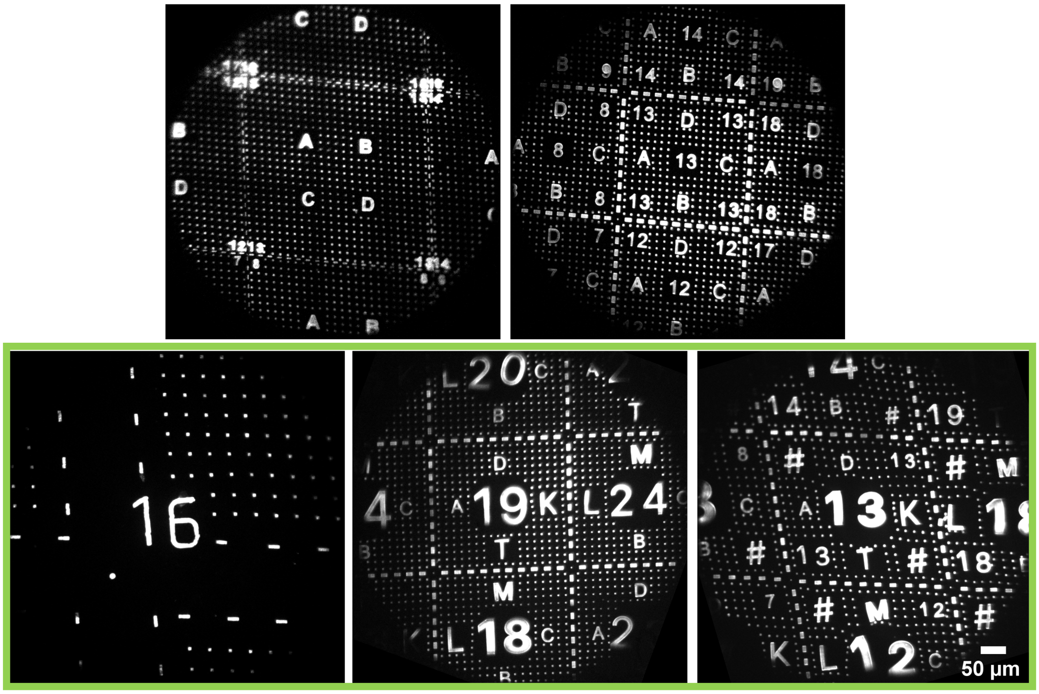


**FIGURE S10** | Optimization of fiducial markers design to enhance feature tracking accuracy. Five characteristics of an interested feature (centroid, filled area, shape, euler number, and perimeter) were extracted using regionprops library for identification of the same features through nearest neighbor algorithm. In order for the algorithm to distinguish and track the correct features, each feature in the field of view has to be unique from its adjacent or nearby features. Hence, some adjustment of the designs and sizes of fiducial markers were done. Five different designs are shown in the figure with green box highlighting the designs that works well for AutoTracking software. Overall, the size of the markers need to be big enough for accurate image binarization and similar feature shape should be avoided.

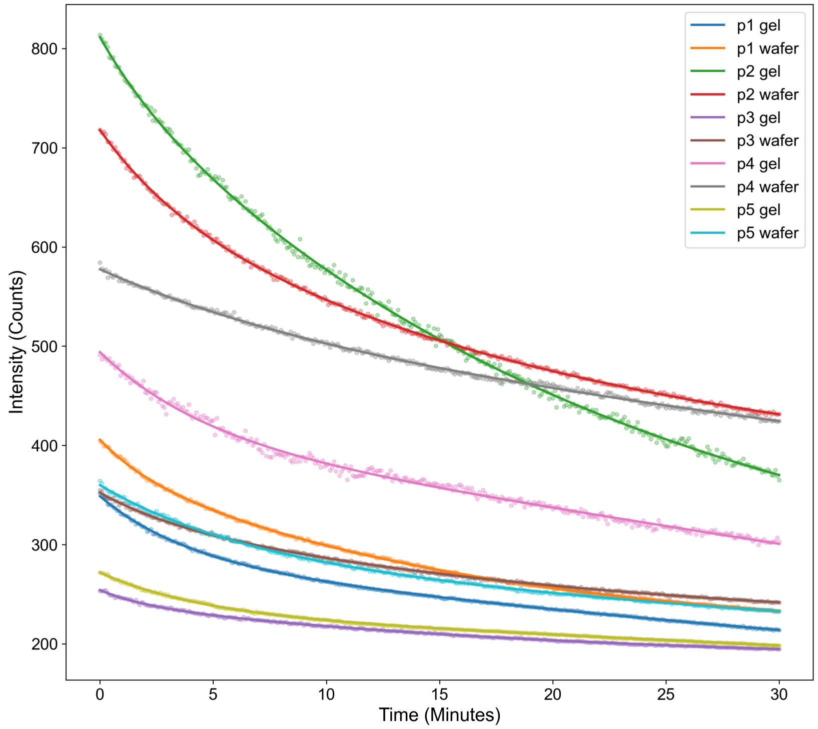


**FIGURE S11** | Photobleaching optimization of imaging parameters used for AutoTracking software. To ensure successful tracking of fluorescence markers using AutoTracking software, power laser (milli-watts) and exposure time (milliseconds) are the two main parameters that need to be optimized. Fluorescence markers are prone to photobleaching after prolonged exposure to laser which later affects intensity counts and results in features with fuzzy edges that can lead to out-of-focus images. The photobleaching experiments were conducted on fiducial markers (Nanoscribe IP-Visio resin) doped with Atto-633 fluorescence dye on both silicon wafer and after transfer into hydrogel. Four variations of imaging conditions were used: (p1) power = 5 mW, exposure time = 10 ms, (p2) power = 5 mW, exposure time = 50 ms, (p3) power = 2.5mW, exposure time = 10 ms, (p4) power = 2.5 mW, exposure time = 50 ms, and (p5) power = 3.5 mW, exposure time = 10 ms. Fiducial markers on both substrates were exposed to 633 nm wavelength laser at the same area for 30 minutes with image acquisition at every 5 seconds. Average intensity of the fiducial markers at each time point were analyzed from a small region of interest in image center then plotted against time. Total laser exposure duration is selected to be 30 minutes to mimic maximum duration when conducted a biological experiment. The results confirm photobleaching effect of fiducial markers regardless of imaging conditions variation. Condition p4 (laser power = 2.5 mW, exposure time = 50 ms) was found to give a successful binarization and accurate AutoTracking result due to the high initial intensity count, slower bleaching rate, and sufficient intensity remained after bleaching. Thus, imaging condition p4 is selected as optimized parameters. We also observed that at similar imaging conditions, fiducial markers on silicon wafer have higher average intensity than markers on hydrogel which may be due to handling and marker transfer process onto the hydrogel.

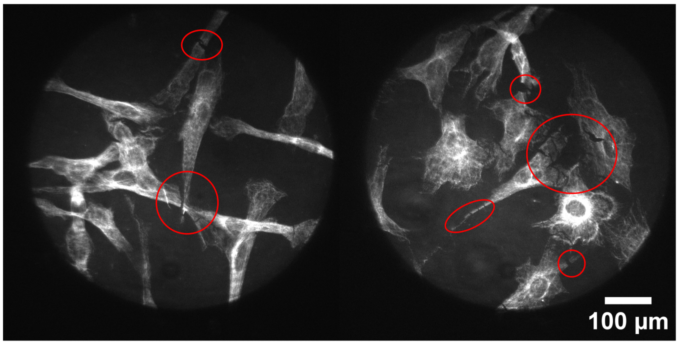


**FIGURE S12** | Monitoring of changes in sample can be done through AutoTracking. Here, we provides an example of fixed NIH 3T3 tagged with Alexa-488 microtubule imaged using 488 nm laser at power density of $5\ mW/cm^2$ and 150 ms exposure time where cell fragmentation is visible during expansion. With the AutoTracking capability, we can identify the exact timing of biological changes.

### 6 | Application to cellular expansion

#### 6.1 | Fixed cells on hydrogel

NIH 3T3 fibroblast cells were cultured in a native manner on ITO-coated coverslip imprinted with fiducial markers in a six well plate. Standard fixation and staining procedures as described in [9] were done prior to treatment with Acryloyl-X (AcX) to cross-link the cells and hydrogel. Hydrogel (DN-Stiff, described in 2) is poured onto the treated cells to allow for polymerization in a sandwich glass mold. Then, the hydrogel was peeled from the ITO glass coverslip with care prior to soak in 20 mM Calcium Sulfate slurry to form Alginate-Calcium ionic crosslinks and complete the double network synthesis. Finally, the fixed cells on hydrogel sample was treated with Proteinase K digestion buffer to mechanically homogenizing the sample, allowing for uniform expansion. The hydrogel was cut to size before loading onto the stretcher.

#### 6.2 | Live cells on hydrogel

HeLa cells were cultured in a six well plate using a complete growth medium DMEM supplemented with 10% FBS, following a standard cell culturing protocol [9]. DN-Medium hydrogel (described in 2) were sterilized by ultraviolet (UV) irradiation [10] using a 254 nm UV source (Amazon). Samples were exposed to UV for 20 min, rested for 10 min, and then re-exposed for an additional 20 min. This sterilization protocol was applied consistently to all hydrogels. Hydrogels were incubated in 3 mL of complete growth medium DMEM supplemented with 10% FBS for 30 min at 37°$C$ and 5% CO2 to equilibrate prior to cell seeding. Growth medium was aspirated and hydrogels were allowed to dry for 30 minutes under sterile conditions before use. HeLa cells were seeded onto the air dried hydrogels at 1 million cells per mL in 250 microlitters of culture medium. Hydrogels were then incubated at 37°$C$ with 5% CO2, and left overnight for cell adhesion. The hydrogel with adhered, live cells is carefully loaded into the iris device with the help of the mounting system. Care is taken to ensure cells are not removed from the hydrogel surface.

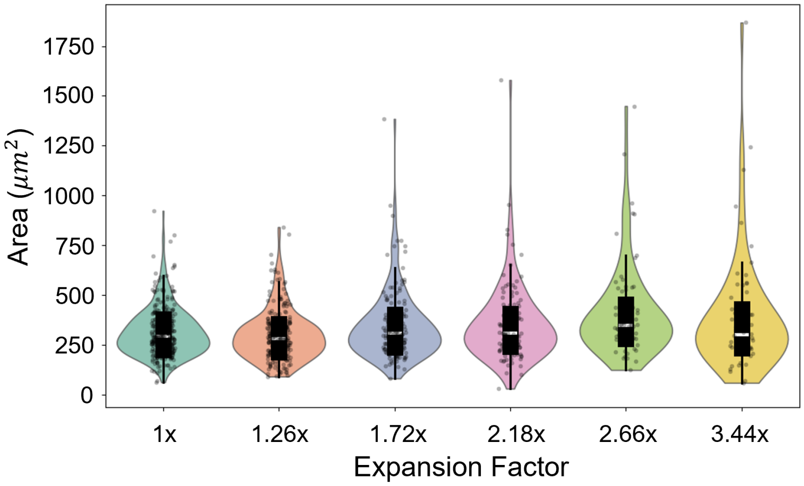


**FIGURE S13** | Analysis of area expansion of live HeLa cells across the full expansion. Post-stretch images at different expansion factor were analyzed for area using cell segmentation software (Cellpose) [11] where false positives were manually removed. The violin plot shows no significant increase in cell area after expansion. On the other hand, the population of HeLa cells (represented as circular dots) significantly decreases after expansion, revealing cell separation since the number of cells decreases when compared in a field of view. Note that the expansion factor shown is scaled based on tautness of hydrogel which limits the full expansion to 3.44$x$.

### 7 | Additional resources

1. Assembly video of the stretcher can be found here: https://www.youtube.com/watch?v=uJTlgtQk6u0
2. Open source CAD design files, assembly manual, AutoTracking code, and fiducial markers real-time tracking videos are uploaded in Kisley lab Github at https://github.com/KisleyLabAtCWRU/TExM.
